# Attributing the temperature response of tree seedling growth to underlying mechanisms

**DOI:** 10.64898/2025.11.29.691274

**Authors:** Dushan P. Kumarathunge, Kashif Mahmud, John E. Drake, Mark G. Tjoelker, Francisco J. Cano, Belinda E. Medlyn

## Abstract

Studies of plant responses to temperature often focus on rates photosynthesis and respiration. However, within-plant utilisation and allocation of carbon are also strongly affected. It is unclear how much each of these processes contribute to determining the overall temperature response of growth. We applied a data assimilation framework to a glasshouse experiment with detailed physiological and growth measurements to investigate the relative contribution of different physiological processes to the overall temperature response of tree seedling growth. We found that both short-term effects of temperature and acclimatory responses of photosynthesis and respiration had a significant impact on the temperature response of growth. However, the effect of temperature on biomass allocation patterns to different tissues, non-structural carbohydrate utilisation and C losses to other unmeasured losses were also substantial in determining the temperature response of growth, particularly at sub-optimal temperatures. Our work demonstrates that the growth response to warming cannot be predicted using only the direct effect of temperature on photosynthesis and respiration and emphasizes the importance of temperature acclimation of photosynthesis, respiration and other C balance processes. Our results provide new guidance for process-based models to correctly describe the temperature effects on tree growth.

## Introduction

The rate of tree growth is strongly determined by the environment. Tree metabolic processes are directly affected by atmospheric CO_2_ concentration, light, temperature, soil water and nutrient availability; significant changes in these drivers can affect tree growth in both positive and negative ways (Lambers *et al*., 2008). Out of these factors, temperature plays a central role because it affects metabolic rates of every physiological process (Kumarathunge et al., 2019; Lu et al. 2013; Way and Oren 2010; Körner, 2008; Atkin et al., 2005). Understanding and quantifying how tree growth is affected by temperature is critically important in predicting the impact of global warming on the terrestrial carbon (C) cycle (Kumarathunge *et al*., 2019, Lombardozzi *et al*., 2015, Rogers *et al*., 2017a, Smith & Dukes, 2013).

Process based vegetation models (PBMs) are the principal tool used to predict the impact of global warming on terrestrial vegetation (Mäkelä *et al*., 2000, Medlyn *et al*., 2011b).

Typically, PBMs represent the growth of forests as the difference between C uptake through photosynthesis, and C loss through respiration and turnover. Hence, the temperature response of tree growth emerging from these models depends strongly on the parameterisations used for photosynthesis and respiration. In many models, the temperature responses of photosynthesis and respiration are assumed to be static (Lombardozzi *et al*., 2015). However, much of the recent physiological literature suggests that both photosynthesis and respiration undergo thermal acclimation (Wu et al., 2025, Rubio et al., 2025, Carter et al., 2024, Choury et al., 2022, Drake *et al*., 2019, Drake *et al*., 2016, Way & Yamori, 2014, Crous *et al*., 2013, Gunderson *et al*., 2009, Hikosaka *et al*., 2006, Atkin & Tjoelker, 2003). There is also increasing awareness that leaf-to-air vapour pressure deficit, which increases with warming, can have a large impact on leaf physiology (Lopez et al., 2021). There is thus a very significant effort to experimentally quantify how leaf physiology responds and acclimates over time to warming, including responses of photosynthesis, stomatal conductance, and respiration, to warmer temperatures higher VPD (Zheng et al., 2025, Cox et al., 2024, Middleby et al., 2024, Cox et al., 2023, Marchin et al., 2023, Choury et al., 2022, Smith & Dukes, 2017, Crous et al., 2013, Gunderson et al., 2009, Hikosaka et al., 2006, Atkin & Tjoelker, 2003). Many PBMs are being updated to include these responses (Knauer et al., 2023, Mengoli et al., 2022; Oliver et al., 2022, Huntingford *et al*., 2017, Smith *et al*., 2017, Smith *et al*., 2016). This focus on net C uptake as the key process of interest is reinforced by the similarity between the temperature response of acclimated photosynthesis and that of growth (Garen et al., 2024).

However, analysis of plant growth responses to temperatures using a traditional relative growth rate (RGR) framework (Fetcher et al., 2022, Drake et al., 2027, Puglielli et al., 2017) typically highlights an important role for other plant processes associated with C utilisation, including changes in specific leaf area (SLA) (Poorter et al., 2012, Poorter et al. 2010, Poorter et al. 2009, Scheepens et al., 2010, Atkin et al., 2009) and shifts in biomass allocation (Blessing et al., 2015, Cheesman & Winter, 2013). For example, in a temperature response experiment with *Eucalyptus tereticornis* seedlings, Drake et al. (2017) found that changes in leaf area ratio (LAR, reflecting C utilisation) were larger and more strongly related to plant growth rate than were changes in net assimilation rate (NAR, reflecting C uptake), despite large changes in photosynthesis and respiration rates. Changes in C utilisation patterns with temperature are commonly not included in PBMs because they are less straightforward to parameterise from experimental studies than are gas exchange data. Nonetheless, their importance is underscored by several modelling studies suggesting that growth responses cannot be predicted from physiological responses alone (Campany *et al*., 2017, Fatichi *et al*., 2014, Körner, 2003, Körner, 2015, Leuzinger *et al*., 2013), indicating that other C balance processes must also be important (Mahmud *et al*., 2018). A comprehensive understanding of how these additional processes contribute to the overall response of plant growth to temperature is needed to accurately predict the impact of warming on plant growth by PBMs. To do so, we need to analyse experimental data with a quantitative framework that can integrate all the individual processes of interest.

Here, we utilised a data assimilation (DA) framework (Williams *et al*., 2005) to infer shifts in C utilisation processes in a warming experiment with tree seedlings. The DA framework involves fitting a simple, minimal-assumption carbon balance model of a plant to all available data, including physiological and growth data, to infer flows of carbon in the plant over time, and hence quantify these hidden processes. Not only does this approach provide a quantitative estimate of allocation parameters, it also allows information on photosynthesis and respiration to be directly linked to the resultant patterns of growth, making it possible to quantify the contribution of individual mechanisms to the overall temperature response of tree seedling growth. Further, it does so in a process-oriented way that can readily be translated to PBMs. DA has been increasingly used to analyse plant C balance processes, including dynamics of non-structural C (Richardson *et al*., 2013), seasonal shifts in biomass allocation and respiration in tree species (Rowland *et al*., 2014), responses of stem growth to water availability in urban forests (Zheng et al., 2023), to resolve environmental limitations on the growing-season onset of gross primary production (Stettz et al., 2022), and to simulate continental-scale patterns in C cycle processes (Bloom *et al*., 2016).

Mahmud et al. (2018) utilised a DA framework to infer the effects of belowground sink limitation on C balance processes of *Eucalyptus tereticornis* seedlings. Here, we apply the approach taken by Mahmud *et al*. (2018) to analyse data of a previously published experiment in which *Eucalyptus tereticornis* seedlings were grown at six growth temperatures in a temperature-controlled glasshouse (Drake et al., 2017). We use a simple scaling approach to estimate total plant C uptake from measurements of leaf photosynthesis and tissue respiration. We then use data assimilation of a simple C balance model to infer fluxes of carbon within the plant, including carbohydrate utilisation and biomass allocation, following Mahmud *et al*., (2018).

These steps are applied at each of the six growth temperatures. We then carried out a sensitivity analysis to quantify the importance of each of the C uptake and utilisation processes in determining the overall temperature response of tree seedling growth. The key insight of this work is that while photosynthesis and respiration matter, they were insufficient to predict growth responses to warming. Below optimal temperatures, changes in specific leaf area, allocation patterns, and respiration acclimation drove most of the growth increase. Above optimal temperatures, direct temperature effects on gas exchange dominate but were modulated by acclimation responses.

## Materials and methods

### Experimental design

We used data from a previously published empirical study (Drake *et al*., 2017) that tested the effects of climate warming on growth and physiology of forest red gum (*Eucalyptus tereticornis*) seedlings. Drake *et al*., (2017) measured a range of physiological and growth variables under six daily mean growth temperature treatments; 18, 21.5, 25, 28.5, 32 and 35.5 °C. A detailed description of the experimental design can be found in Methods S1 and Drake *et al*. (2017).

### Carbon balance model (CBM)

We used a simple minimal-assumption C balance model (Figure 1) in this study following Mahmud *et al*., (2018). Models of this form often form the mechanistic core of PBM across scales from ecosystems to global models. Our model was driven by daily gross primary production (*GPP*) inputs. The daily maintenance respiration (*R*_m_) was subtracted from *GPP* and the remaining C entered a non-structural C pool (*C_n_*). C stored in the *C_n_* pool was utilised for growth at a rate *k*. A fraction of the remaining C was allocated to structural C pools in leaves (*C_s,f_*), wood (*C_s,w_*) and roots (*C_s,r_*). The fraction *Y* was the unmeasured C losses by the plant which includes fluxes such as growth respiration, root exudation and volatile organic C emission. We did not consider turnover of leaves, wood or roots as the seedlings were young, growing rapidly and the total duration of this experiment was short (∼48 days). We quantified the dynamics of each C pool using four equations (Eqn 1- 4).

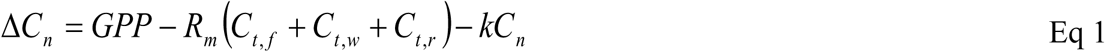

**Figure 1.**
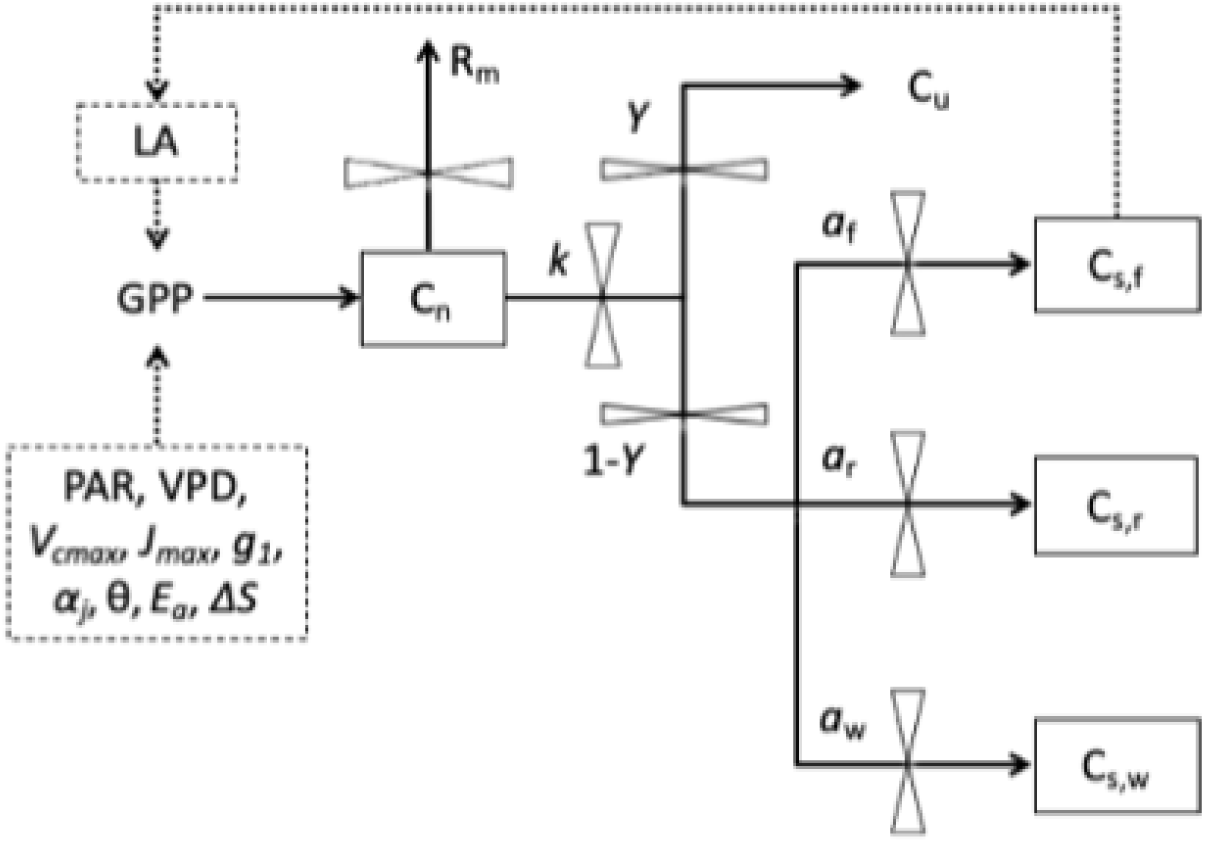
Structure of the Carbon Balance Model used in this study. Arrows depict the C fluxes: GPP, gross primary production; PAR, photosynthetically active radiation; VPD, vapor pressure deficit; *V*_cmax_, maximum rate of ribulose-1,5-bisphosphate carboxylase-oxygenase (Rubisco) activity; *J*_max_, potential rate of electron transport; *g*_1_, stomatal conductance parameter; *E_a_*, activation energy of *V*_cmax_ and *J*_max_; *ΔS*, entropy of *V*_cmax_ and *J*_max_, *α_J_*, quantum yield of electron transport; θ, the curvature of the light response of electron transport rate; *R*_m_, total measured respiration; *C*_u_, unmeasured C losses. Boxes depict the C pools: *C*_n_, non-structural storage C; *C*_s,f_, structural C in foliage; *C*_s,r_, structural C in roots; *C*_s,w_, structural C in wood. Fluxes are governed by five key parameters: *k*, storage utilization coefficient; *Y*, growth respiration and other unmeasured C losses; *a*_f_, allocation to foliage; *a*_w_, allocation to wood; *a*_r_, allocation to roots.

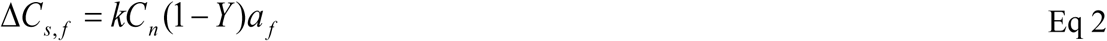

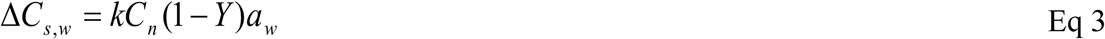

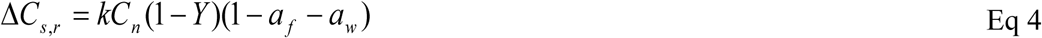

where *GPP* is the gross primary production (gC plant^-1^ d^-1^), *R*_m_ is the measured maintenance respiration rate (gC g^-1^C d^-1^), *C*_t,f_, *C*_t,w_, and *C*_t,r_ are the total C in foliage, wood and root respectively (g C plant^-1^), *k* is the storage utilization coefficient (g C g^-1^ C d^-1^); *Y* is the fraction of unmeasured C lost (this term includes growth respiration and other unmeasured C losses such as root exudation and volatile organic C emission), and *a_f_, a_w_,* and *a_r_* (i.e. 1-*a_f_-a_w_*) are the allocation to foliage, wood and root respectively.

The nonstructural C pool (*C_n_*) is assumed to be divided amongst foliage, wood and root tissues (*C*_n,f_, *C*_n,w_, *C*_n,r_) with a ratio of 75 : 16 : 9, utilizing data from an experiment on 4-month-old *E. globulus* seedlings (Duan et al., 2013). Both nonstructural C (*C_n_*) and structural C (*C_s_*) in each tissue was then summed up to estimate the total C in that tissue (*C_t_*).

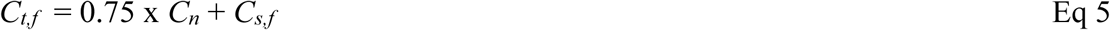

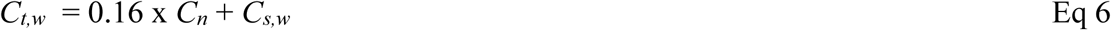

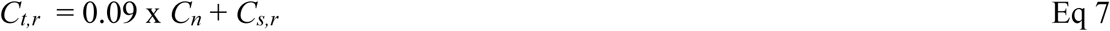

We used DA to estimate the parameters of the C balance model (*k, Y, a*_f_*, a*_w_*, a*_r_). Daily total GPP and *R_m_* were inputs to the DA framework, and the measurements of total C mass of each of the plant (48% of the total biomass) components were used to constrain the parameter values. Our DA approach used the Metropolis algorithm as implemented by Zobitz *et al*. (2011). The Metropolis algorithm (Metropolis et al., 1953) is a Markov Chain Monte Carlo method for acquiring a sequence of random samples from a probability distribution from which direct sampling is challenging. To account for potential changes over time (e.g., ontogeny) all parameters were represented as a linear function of time, *t* (*p* = *p*_1_ + *p*_2_*t*), hence a total of eight parameters were fitted. A detailed description of the methods of the DA framework is given by Mahmud *et al*. (2018).

We used broad uniform prior probability density functions for *k, a*_f_, and *a*_w_ (Table 1). We constrained the probability density function of the parameter *Y,* based on the literature on growth respiration (Amthor, 2000). We accounted for the uncertainty in DA outputs by considering standard errors of input variables using multiple replicate measurements for *GPP*, *R_m_* and C mass of each of the plant components.

**Table 1.** Prior parameter probability density functions and their starting values that were used in DA to estimate the parameters of the C balance model.

| Parameter | Minimum | Maximum | Starting Value |
| --- | --- | --- | --- |
| $k$ | 0 | 1 | 0.5 |
| $Y$ | 0.2 | 0.4 | 0.3 |
| $a_f$ | 0 | 1 | 0.5 |
| $a_w$ | 0 | 1 | 0.5 |
| $a_r = 1 - (a_f + a_w)$ | | | |

We tested the importance of a non-structural C storage pool in the C balance model by comparing the full model (Figure 1) with a simplified model without the non-structural C pool.

We performed DA with both models and the model performance was compared using the Bayesian Information Criterion (BIC; Schwarz, 1978).

### Estimation of GPP

The daily total GPP of seedlings was calculated using Eqn 10.

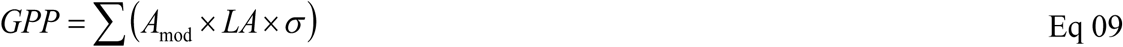

where, *A*_mod_ is the modelled leaf net photosynthesis rate, LA is total seedling leaf area (m^2^) and σ is the self-shading factor (Campany *et al*., (2017). We estimated the self-shading parameter for each growth temperature treatment by using 61 pre-digitized Eucalyptus seedlings from Duursma et al. (2012). This approach used a 3D plant structure built using digitized plant allometry and the photosynthetic parameters obtained in the experiment to predict photosynthesis using total seedling leaf area. The self-shading factor for each temperature treatment was calculated as the ratio of total photosynthesis with self-shading to a sun lit horizontal leaf. We then developed a simple linear regression model to estimate self-shading factor as a function of seedling total leaf area using the data from 61 digitised seedlings. The self-shading factor for each growth temperature treatment was predicted for each day using the previous days’ cumulative leaf area and ranged from 0.88 – 0.90.

We used a coupled photosynthesis-stomatal conductance model to quantify leaf net photosynthesis rates (*A*_mod_) of seedlings at each growth temperature. The standard biochemical model of photosynthesis (Farquhar *et al*., 1980) was coupled to the optimal stomatal conductance model proposed by Medlyn *et al*. (2011a). The model was run with photosynthetic parameters, namely the maximum rate of ribulose-1,5-bisphosphate carboxylase-oxygenase (Rubisco) activity (*V*_cmax_), temperature response parameters of *V*_cmax_ and *J*_max_, stomatal conductance model parameter (*g_1_*), and the potential rate of electron transport (*J*_max_), quantum yield of electron transport (*αj*) and the curvature of the light response of electron transport rate (*θ*)) and meteorological data namely PAR, air temperature and VPD in the glasshouse. We estimated *A*_mod_ for each growth temperature treatment at 15-minute intervals by driving the leaf gas exchange model (*Photosyn* function within plantecophys R package, Duursma, 2015). We scaled these leaf-level values to the whole plant by multiplying seedling total leaf area and a self-shading factor, following Campany *et al*. (2017). The daily total gross primary productivity (*GPP*_total_) for each plant was then calculated by converting estimated photosynthetic rates to mass C gain over 15-min time steps and summing for 24 h. A detailed description on the measurement protocols used to collect gas exchange data and estimation of photosynthetic parameters are given in methods S1.

### Estimation of respiration (R_m_)

We estimated tissue-specific *R*_m_ for each growth temperature treatment at 15-minute intervals. We used the measured tissue-specific respiration rates at a common temperature of 25 °C to estimate total seedling respiration in each growth temperature, assuming all tissues share a common short-term temperature response as measured for leaves (Eqn 11).

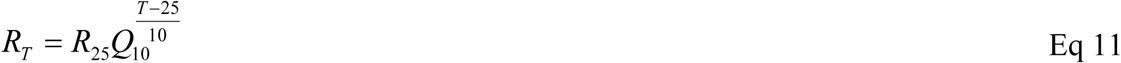

where *R_T_* is the tissue specific rate of *R*_m_ at temperature *T*, *R*_25_ is the rate of *R*_m_ at a reference temperature of 25 °C, and *Q*_10_ is the multiplicative rate of increase in *R*_m_ for every 10 °C change in temperature. The daily *R*_m_ for each plant was then calculated by converting estimated total leaf, wood, and root *R*_m_ to mass C loss over 15-min time steps and summing for 24 h. A detailed description of the measurement protocols used to collect respiration data and estimation of respiratory parameters are given in methods S1.

### Sensitivity analysis

We performed a sensitivity analysis to quantify how individual processes (Figure 1, Table 2) contribute to overall plant growth by the sequential addition of a single process at a time. In this analysis, we calculated the daily total GPP using the modelled leaf area to account for leaf area feedback through its effect on foliage mass, and consequently GPP, over time following the procedure of Mahmud *et al*., (2018). Modelled leaf area is calculated by multiplying modelled foliage mass by a time series of specific leaf area (SLA). Firstly, we ran the C balance model with the inputs and modelled parameters from the lowest growth temperature (18 °C), then changed the parameters to those for the optimum temperature for seedling growth in this experiment (28.5 °C). We sequentially changed the modelled parameters one at a time to quantify the effect of each parameter on the response to increasing temperature from below optimum to the optimum. For example, attribution Case 1 considers the baseline parameter setting of 18 °C and the direct temperature response of photosynthesis, while attribution Case 2 has the same settings plus the direct temperature response of respiration. This sequential approach allows us to test what responses are necessary in a model to correctly represent the overall effects of temperature on growth: for example, is it sufficient to represent direct responses only, or direct responses plus temperature acclimation? We repeated the same procedure with the inputs and modelled parameters from the optimum temperature for growth (28.5 °C) to the highest growth temperature (35.5 °C), quantifying the effect of each parameter on the final seedling biomass at growth temperatures above the optimum. The parameters considered for the sensitivity analysis are given in Table 2.

**Table 2.**
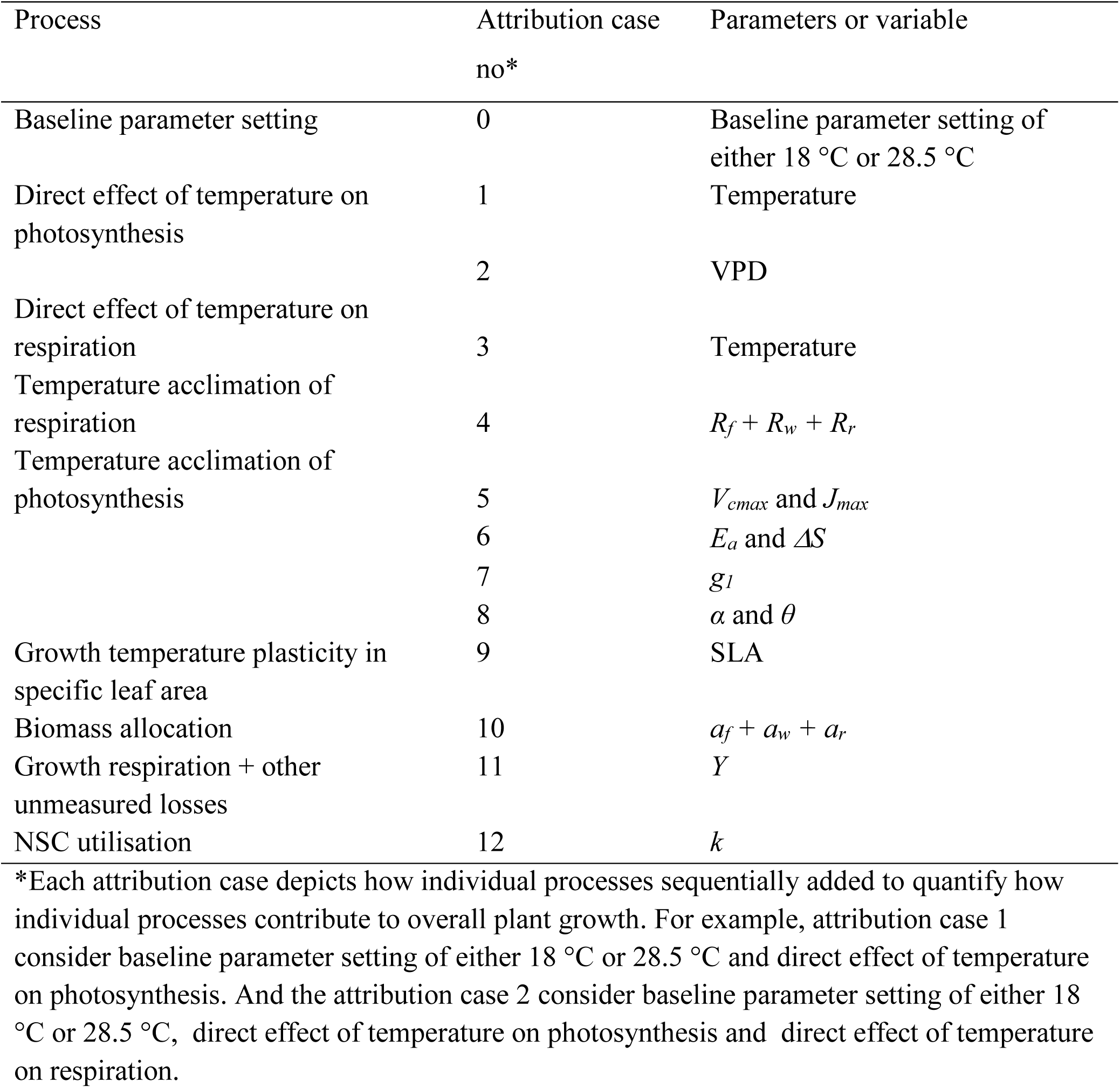
Different processes and their parameterization included in the sensitivity analysis to quantify how individual processes contribute to overall plant growth.

## Results

### Daily carbon gain

Modelled daily photosynthetic rate per unit leaf area (*GPP*; gC m^-2^ d^-1^) diverged between temperature treatments from the start of the experiment (Figure. 2a). *GPP* was lowest at the highest growth temperature throughout the experiment. The highest *GPP* values were observed for seedlings grown at intermediate growth temperatures, 21.5 and 28.5 °C, up to 6 weeks after the start of the experiment. By the time of the final harvest, *GPP* was highest at the coolest growth temperature (18 °C) and lowest at the warmest growth temperature (35.5 °C).

**Figure 2.**
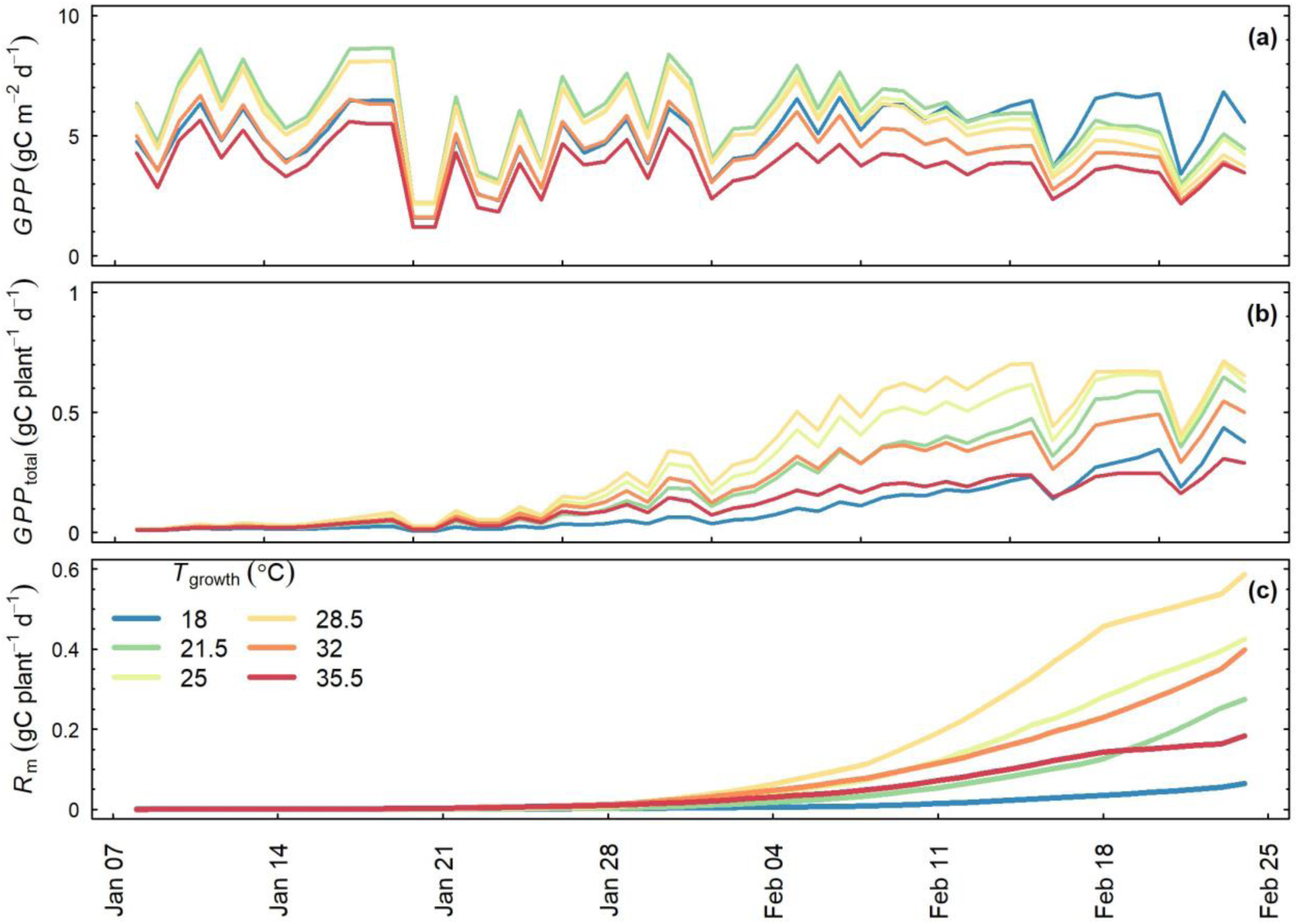
Time series of the C fluxes at six growth temperatures (*T*_growth_): (a) modelled daily photosynthetic rate per unit leaf area, (b) daily total GPP and (c) daily total respiration (*R*_m_). Major ticks on the x-axis reflect start of the weeks.

As leaf area increased over time, the daily total gross primary productivity (*GPP*_total_) showed an exponential increase with time (Figure. 2b). At the beginning of the experiment, the lowest *GPP*_total_ was estimated for seedlings at 18 °C while the highest was for plants at 28.5°C. However, at the end of the experiment, the 35.5 °C-grown seedlings showed the lowest *GPP*_total_.

### Daily respiration (R_m_)

Similar to *GPP*_total_, total *R_m_* showed an exponential increase with time (Figure.2c), reflecting the approximately exponential increase in biomass over time (Drake et al., 2017). Daily total *R_m_* was highest for the seedlings grown at 28.5 °C and lowest for those grown at 18 °C. The largest contribution to whole-plant respiration rates came from leaves, reflecting the fact that they formed the largest biomass pool (Figure S5).

### Carbon stock dynamics

The C balance model was able to reproduce the tissue-specific biomass growth over time in response to growth temperature treatment (Figure 3). Most of the predicted biomass values were within one standard error of the measurements except for the final measurement where the predicted values were slightly higher compared to the measurements. Leaf, wood and root mass diverged between the growth temperature treatments approximately two weeks after the start of the experiment. Both the lowest (18 °C) and the highest (35.5 °C) growth temperatures showed significantly lower leaf mass compared to the other growth temperatures (Figure 3a). At the end of the experiment, wood mass of 18 °C-grown seedlings was significantly lower compared to the other treatments (Figure 3b) while the highest wood mass was found at 28.5 °C. Similar to leaf mass, both the lowest (18 °C) and the highest (35.5 °C) growth temperatures showed significantly lower root mass compared to the other growth temperatures (Fig. 3c). The modelled total non-structural carbohydrate pool (gC plant^-1^) differed among growth temperature treatments; a lower non-structural carbohydrate pool was inferred for both the lowest (18 °C) and the highest (35.5 °C) growth temperatures compared to the other growth temperatures (Figure 3d).

**Figure 3.**
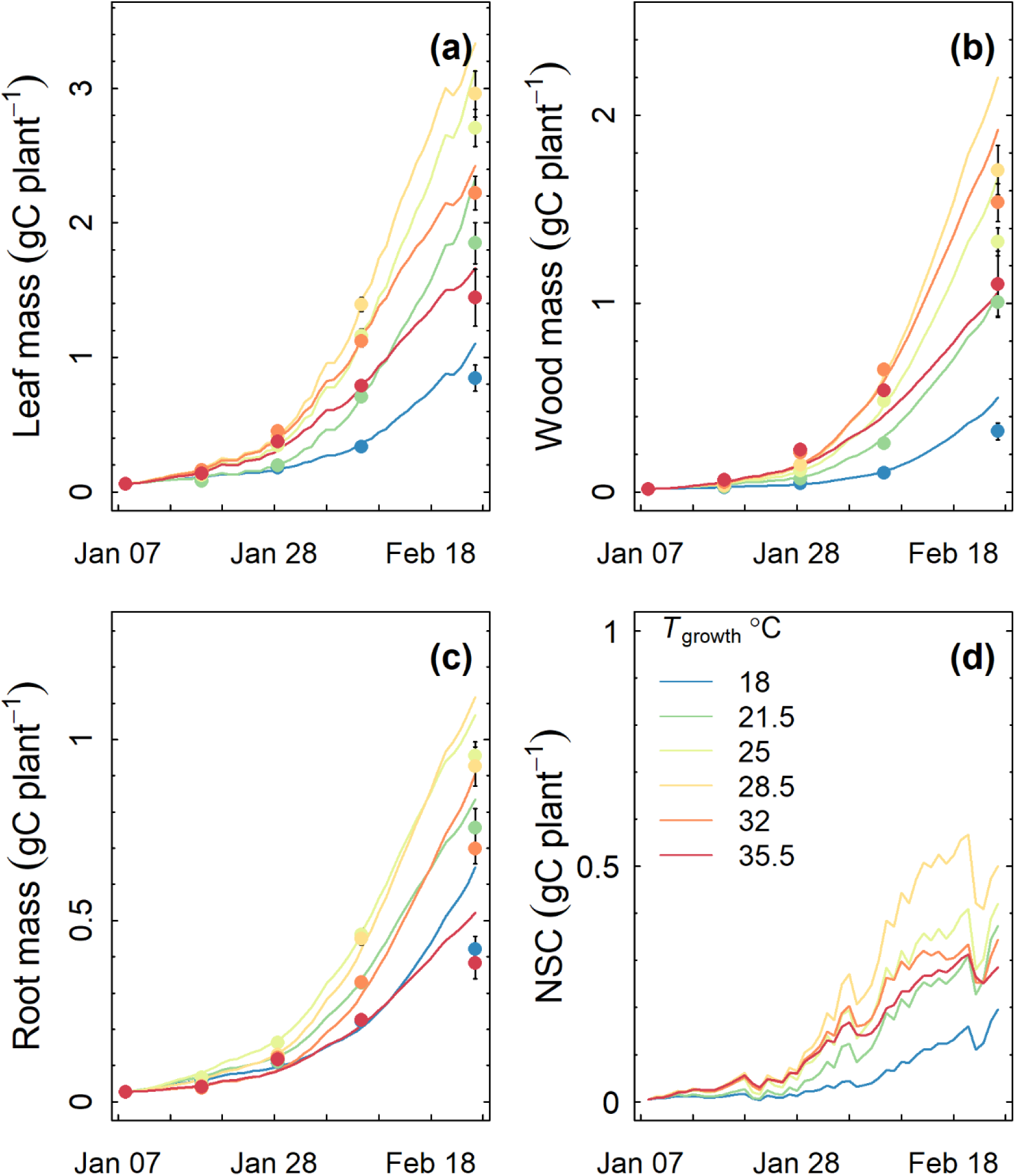
Time-series of daily modelled C stocks (lines) at six growth temperatures (*T*_growth_) with corresponding measured data (filled circles): (a) total C mass in foliage, (b) total C mass in wood, (c) total C mass in root and (d) total C mass in non-structural carbohydrate pool (C_n_, See Figure 1). Error bars depict 1SE. Major ticks on the x-axis reflect weeks.

### Parameters of the carbon balance model

We inferred growth temperature effects on all five fitted parameters of the C balance model (Figure. 4). When averaged across time, both the utilization coefficient (*k*) and unmeasured C losses by the plant (including growth respiration and other unquantified losses such as root exudates and volatile organic C emission; *Y*) showed a peaked response, with the maximum rates being estimated at intermediate growth temperatures (21.5 – 28.5 °C; Figure 4a, b). The utilization coefficient (*k*) showed a decreasing trend as the experiment progressed (Figure S6a). The rate of the decline in *k* with time was larger at both the coldest and warmer growth temperatures compared to the intermediate growth temperatures, causing a build-up of C storage towards the end of the experiment particularly at higher growth temperatures. In contrast to *k*, *Y* did not vary significantly over time (Figure S6b).

**Figure 4.**
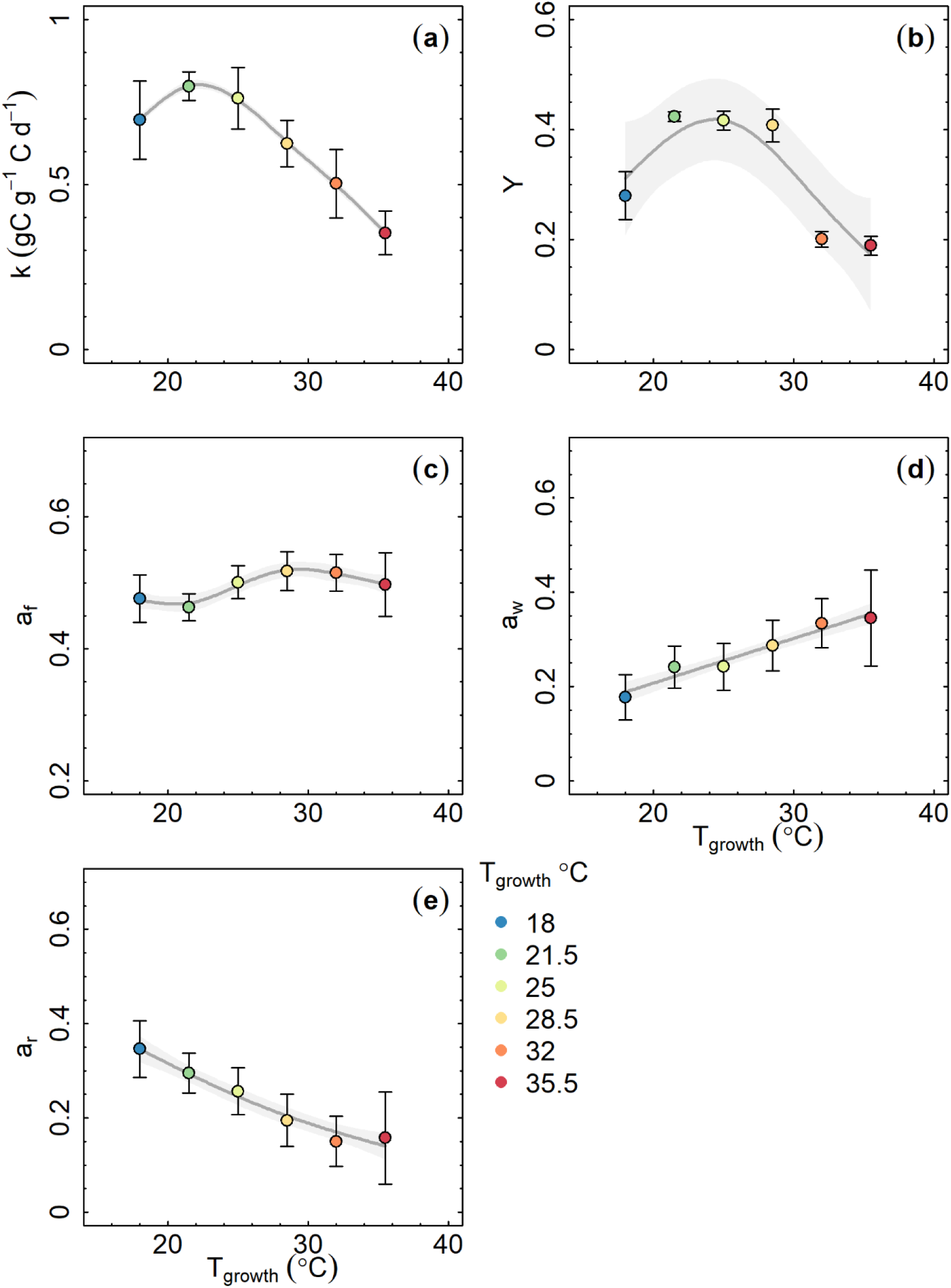
Temperature response of the parameters of C balance model: (a) storage utilization coefficient, *k*; (b) growth respiration and other C losses fraction, *Y*; (c) allocation to foliage, *a_f_*; (d) allocation to wood, *a_w_*; (e) allocation to roots, *a_r_*. The circles represent the average value of the modelled parameters over the entire experiment period (using Figure S5). Lines depict the fitted generalized additive model (GAM) with shaded area showing the 95% CI of predictions. Error bars depicts ±1SE.

The DA process showed a clear divergence in biomass allocation to different tissues at different growth temperatures and over time. Initially, the fractional biomass allocation to foliage (*a_f_*) was highest at 28.5 °C (0.7) compared to the mean *a_f_* at other growth temperatures (0.5, Figure S6c). However, *a_f_* decreased with time at temperatures above 25 °C in contrast to the increasing trend observed at 18 and 21.5 °C growth temperatures. At the end of the experiment, *a_f_* was significantly higher (non-overlapping 95% CIs) at 18 °C (0.5) compared to the mean *a_f_* at higher growth temperatures (0.4). When averaged across the experimental period, *a_f_* showed relatively little change with growth temperature (Figure 4c).

Allocation to wood (*a_w_*) showed a clear increasing trend with growth temperature (Figure 4d) and showed an increasing trend with time (Figure S5d). In contrast to *a_w_*, the fraction of biomass allocated to roots (*a_r_*) showed a decreasing trend with increasing growth temperature (Figure 4e). *a_r_* was significantly higher at the lowest growth temperature during the initial growth period (Figure S6e). Over time, *a_r_* decreased at 18 – 25 °C growth temperatures while a slight increasing trend was observed at temperatures above 25°C.

### Importance of non-structural carbohydrate pool

Table 3 shows the BIC statistic for model fits; (1) with and (2) without the non-structural carbohydrate pool in the C balance model. For all growth temperature treatments, BIC values were consistently lower for the model including the storage pool compared to the model without it. Although some PBMs assume that each day’s photosynthate becomes structural biomass the following day, this finding demonstrates that including a non-structural carbohydrate storage pool in the C balance model enables a better representation of the observed experimental data.

**Table 3.**
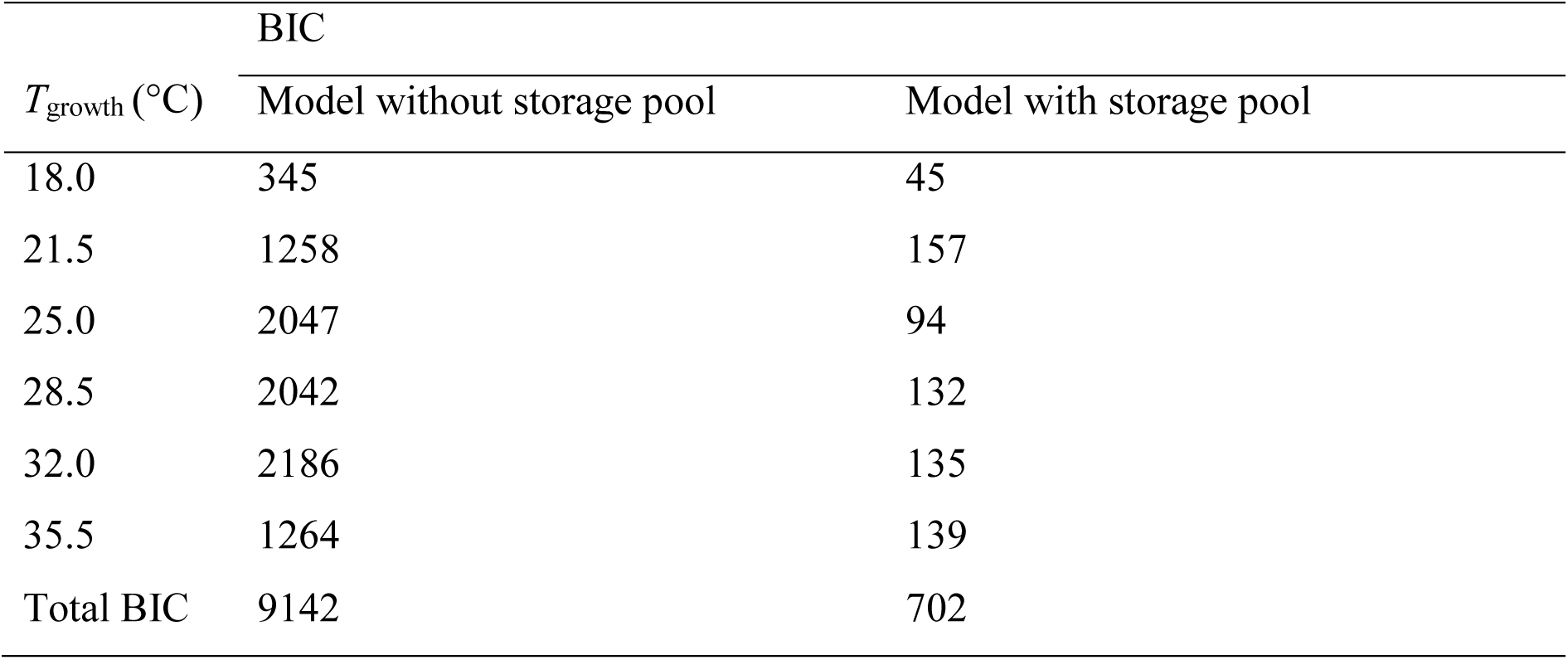
Bayesian Information Criterion (BIC) values for model fits; (1) without and (2) with the non-structural carbohydrate pool in the C balance model.

| $T_{\text{growth}}$ (°C) | BIC | |
| --- | --- | --- |
|  | Model without storage pool | Model with storage pool |
| 18.0 | 345 | 45 |
| 21.5 | 1258 | 157 |
| 25.0 | 2047 | 94 |
| 28.5 | 2042 | 132 |
| 32.0 | 2186 | 135 |
| 35.5 | 1264 | 139 |
| Total BIC | 9142 | 702 |

### Sensitivity analysis

We have shown that temperature affected seedling growth via a range of processes, including short and acclimatory effects on photosynthesis and respiration (see Notes S1 for a detailed results), as well as changes in the non-structural carbohydrate utilization rate, unmeasured C losses, and C allocation to foliage, wood and root pools. We quantified the individual contribution of each of these process responses by running the C balance model with parameter inputs changing one at a time, starting with all parameters set to their value for one temperature and modifying the parameters one by one until all are set to their value for the second temperature (Figure 5). We applied this sensitivity analysis to investigate the change in biomass between i) the lowest growth temperature (18 °C) and the optimum temperature for growth (28.5 °C), and ii) between the optimum temperature for growth and the highest growth temperature (35.5 °C). This provides the key mechanistic insight regarding the processes that are most important to determining tree growth responses to temperature (Figure 5). Given the complexity of these results, we introduce them sequentially in the subsequent sections.

**Figure 5.**
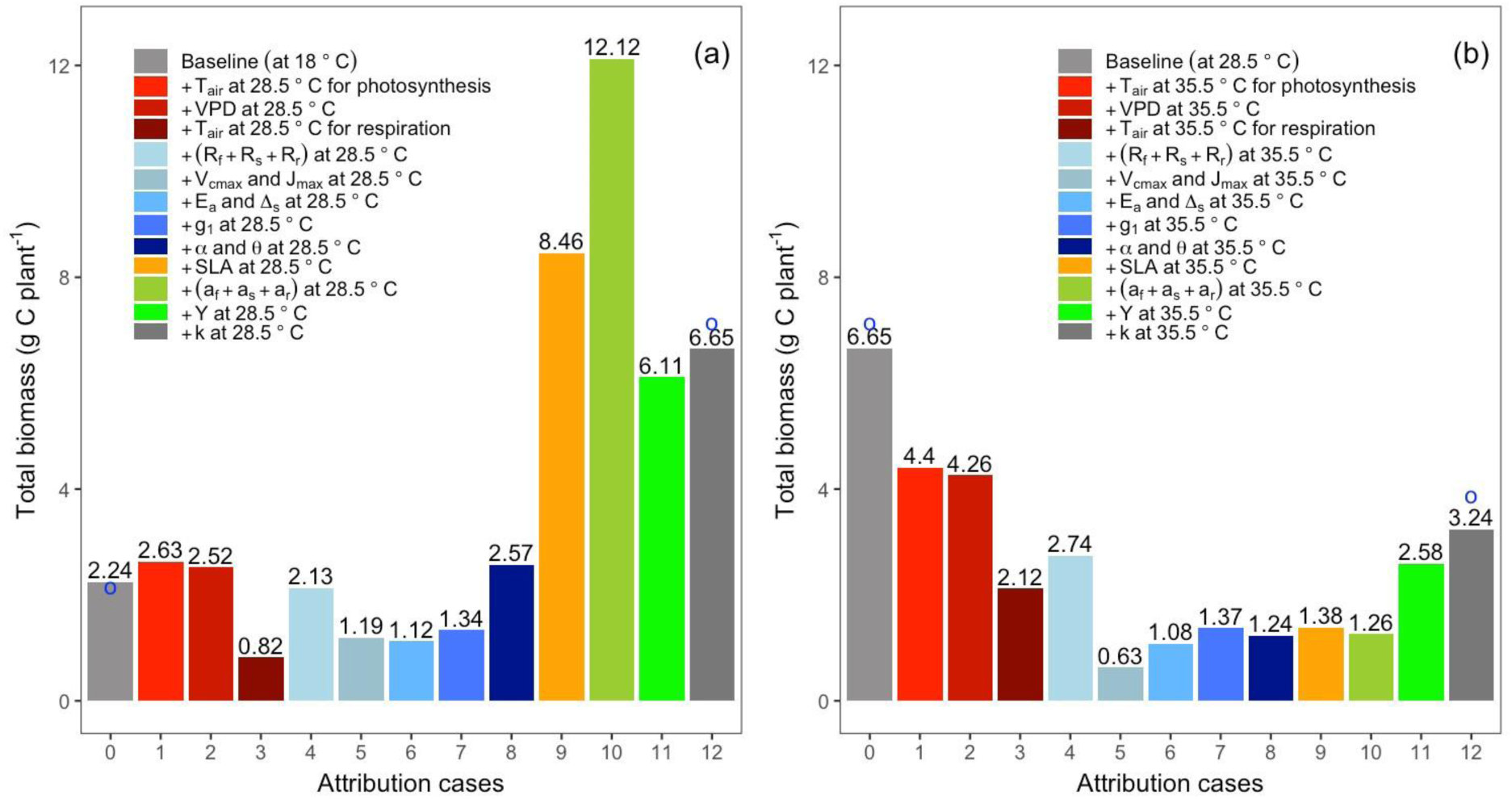
Impact of sequential parameter changes from (a) coolest growth temperature to optimum temperature for growth (18 °C to 28.5 °C) and (b) from optimum temperature for growth to warmest growth temperature (28.5 °C to 35.5 °C) on modelled final total plant mass. Red shades indicate parameters describing direct effects of temperature and VPD (cases 1-3); blue shades indicate parameters describing physiological acclimation (cases 4-8); orange shows specific leaf area (case 9); and green shades show fitted growth parameters (cases 10-11). End cases in each panel (grey; cases 0, 12) show model runs with the full suite of parameters for the growth temperatures investigated. Dots show corresponding measured biomass.

### From low to optimal temperature

Figure 5a shows how modelled total seedling biomass (*C*_t,f_ + *C_t_,*_w_ + *C_t_,*_r_) was affected by each individual process when growth temperature was increased from low (18 °C) to optimal (28.5 °C). Between the lowest and optimal temperatures, observed biomass increased threefold, from 2.24 to 6.65 gC plant^-1^. The direct effects of increased temperature on photosynthesis and respiration could not explain this change. Modelled total biomass was increased by only ∼17% by the direct effect of increasing temperature on photosynthesis (Figure 5a; case 0 vs 1). Adding the effect of increasing VPD on stomatal conductance had a slightly negative effect on the total biomass (∼4% decrease; case 1 vs 2). The small positive impact of increasing temperature on photosynthesis was greatly outweighed by the negative effect of increased respiration at growth temperatures, with the modelled total biomass decreased by 67% (Figure 5a; case 2 vs 3). Thus, a model incorporating only the direct effects of warming on photosynthesis and respiration would predict a decline, not an increase, in growth, at these temperatures (Figure 5a, case 0 vs case 3).

Incorporating temperature acclimation of respiration and photosynthesis into the model did not enable correct prediction of final biomass at optimal temperature. Adding temperature acclimation of tissue respiration (along with the direct effects of warming on photosynthesis and respiration) had a positive effect on modelled growth, but the model still predicted an overall decrease in biomass (Figure 5a, case 3 vs 4). Temperature acclimation of the photosynthetic biochemical capacity, represented by parameters *V*_cmax25_ and *J*_max25_, had a sizeable negative effect on growth, with the predicted final biomass decreasing by approximately 44% between sub-optimal and optimal temperatures (Figure 5a, case 4 vs 5). The photosynthetic temperature response parameters *E*_a_ and *ΔS* did not vary significantly between 18 and 28.5 °C growth temperatures (Table S1) and consequently, their impact on final biomass was small. The stomatal conductance parameter, *g*_1_ varied considerably across treatments (Table S1), but the impact on final biomass was modest; all values of *g*_1_ were high and thus stomatal closure did not restrict C uptake by much. The light-response parameters *α_j_* and *θ* also varied, but when all aspects of temperature acclimation of photosynthesis were taken into account the predicted final biomass at the optimal temperature was not much higher than at the sub-optimal temperature (Figure 5a, case 0 vs case 8).

The largest changes in plant growth modelled at these temperatures were due to changes in specific leaf area (SLA), allocation patterns and growth respiration. SLA increased by 19% from 18 to 28.5 °C (Figure S7), allowing an increase in leaf area for a given leaf mass, and resulting in increased C uptake and growth. Consequently, the change in SLA had a sizeable impact in our sensitivity analysis, increasing final total biomass between sub-optimal and optimal temperature by 5.9 gC (Figure 5a; case 8 vs 9).

The change in inferred allocation fractions (*a_f_*, *a_w_* and *a_r_*) also had sizeable effects on final biomass. The change in allocation parameters from sub-optimal to optimal temperature caused an increase in total biomass by 3.7 gC. The inferred unmeasured C losses were significantly higher at optimum than low temperature (Figure 4b). The effect of this change was to decrease the total biomass by 6.0 gC. The inferred non-structural carbohydrate utilisation rate (*k*) was not significantly different at 28.5 °C compared to 18 °C, therefore the added change was small when going from sub-optimal to optimal growth temperatures (biomass increased by ∼9%).

### From optimal to high temperature

Figure 5b shows how modelled total seedling biomass is affected by each individual process when growth temperature is increased from optimal (28.5 °C) to supra-optimal (35.5 °C). Between the optimal and highest temperatures, observed biomass approximately halved, from 6.65 to 3.24 g plant^-1^ (Figure 5b, case 0 vs case 12). In contrast to the sub-optimal temperatures, the direct effects of temperature on photosynthesis and respiration were large when going from optimal to supra-optimal growth temperatures. The combined effect of increased VPD and temperature on photosynthesis was a reduction in final total mass from 6.6 gC (at 28 °C) to 4.3 gC (at 35.5 °C). The direct effect of increasing respiration further reduced the predicted final biomass to 2.12 gC plant^-1^. This reduction is of the same magnitude as the observed effect of temperature on biomass, suggesting that the reduction in biomass could be predicted from direct effects of temperature alone, but the sensitivity analysis indicates that other processes are also important to consider.

Acclimation of respiration at high temperatures was not as strong as at lower temperatures. Acclimation attenuated the direct effect of respiration somewhat, but not completely: the predicted final biomass including respiration acclimation was 2.74 gC plant^-1^. Temperature acclimation of photosynthesis had a strongly negative effect on the predicted final biomass, which was largely due to the strong reduction in the photosynthetic capacity parameters, *V*_cmax25_ and *J*_max25._ Adjustments in these parameters caused a reduction in predicted total biomass of 77% between optimum and supra-optimum temperatures (Fig. 5b; case 4 vs 5). The observed change in the photosynthetic temperature response parameters *E*_a_ and *ΔS* mitigated this decrease somewhat. When acclimation of all photosynthetic parameters was considered, the final predicted biomass was 1.24 gC plant^-1^, considerably below the observed value (Figure 5b; case 6 vs 12).

Changes in the SLA and allocation had little effect on the predicted final biomass above the optimum temperature. The increase in SLA from optimal to supra optimal temperatures was relatively small (5%, Figure S8), and therefore had a marginal effect on final mass between optimal and supra-optimal temperature (Figure 5b; case 8 vs 9)

The inferred allocation fractions implied a sizeable shift from root allocation to wood allocation (Figure 4d, e), resulting in a larger plant above-ground but a negligible change in total biomass. The inferred unmeasured C losses were significantly reduced at high temperatures (Figure 5b), which had a large effect on predicted final biomass. The inferred non-structural carbohydrate utilisation rate (*k*) also increased, which caused a significant (20%) further increase in the predicted final biomass (Figure 5b; case 11 vs case 12).

Overall, the sensitivity analysis suggested that the increase in growth between sub-optimal and optimal temperatures was principally due to the increase in SLA, temperature acclimation of respiration, and a change in allocation pattern; in contrast, temperature acclimation of *V*_cmax_ and *J*_max_ and the increase in *Y* tended to counteract the increase. The decrease in growth from optimal to supra-optimal temperature was principally due to the short-term direct effects of temperature and VPD on leaf photosynthesis and tissue respiration, temperature acclimation of *V*_cmax_ and *J*_max_, temperature acclimation of respiration and a change in allocation pattern.

## Discussion

In this study, we use a data assimilation framework to quantify the contribution of individual mechanisms towards the overall temperature response of tree seedling growth. Our key finding is that a range of processes contribute to determining the overall response, with different processes being important at sub-optimal temperatures than supra-optimal temperatures. Direct short-term effects of temperature and acclimatory responses of photosynthesis and respiration have a significant impact on the temperature response of growth, but particularly at supra-optimal temperatures. The effect of temperature on non-structural carbohydrate utilisation and C losses to growth respiration and other unmeasured losses were substantial; both of these processes showed a peaked response in which the rates increased up to the optimum temperature for growth (*c*. 28 °C) and decreased thereafter. We also found strong effects of temperature on biomass allocation patterns to different tissues, and these effects were particularly important at sub-optimal temperatures. Overall, we demonstrated that not only photosynthesis and respiration, but also other C balance processes such as the rate of non-structural C utilisation, growth respiration and biomass allocation, and their temperature responses are important in determining the temperature response of tree seedling growth.

### Can we simulate growth responses to temperature through the impact of temperature on photosynthesis and respiration?

Classical approaches in tree growth modelling are often driven by photosynthesis and respiration (Mäkelä et al., 2000; Körner, 2006; Clark et al., 2011; Guillemot et al., 2015; McMurtrie et al., 1990). Hence, in these models, the impact of temperature on tree growth is determined mostly by the direct effects of temperature on photosynthesis and respiration (Dewar et al., 1999; Mcmurtrie & Wang., 1993; McMurtrie et al., 1990). However, our results clearly showed that effect of temperature on growth could not be predicted from the direct temperature and VPD effects on photosynthesis and respiration alone. These effects would, in isolation, imply a much lower optimal temperature for growth than was observed. Although the photosynthetic rates increased with temperature at temperatures below the optimum, this effect was outweighed by the exponential increase in tissue respiration rates at these temperatures. In addition, VPD increases monotonically with growth temperature, and therefore negatively affects stomatal conductance (Middleby et al., 2024, Slot et al., 2024). A PBM incorporating only these effects would predict negative impacts of warming on tree growth across a wide range of temperatures.

### Acclimatory effects of temperature on photosynthesis and respiration

Acclimation of plant respiration to temperature is commonly observed (Aspinwall *et al*., 2016, Drake *et al*., 2016, Heskel *et al*., 2016, Vanderwel *et al*., 2015, Crous *et al*., 2011, Atkin *et al*., 2000, Tjoelker *et al*., 2009, Tjoelker *et al*., 1999), including temperature acclimation of above-ground respiration of trees to warming in field conditions (Drake *et al*., 2016; Drake *et al*., 2019a). The temperature acclimation of tissue specific respiration rates that we observed are in line with patterns observed in past studies (Kumarathunge *et al*., 2020, Crous *et al*., 2017, Drake *et al*., 2017, Atkin *et al*., 2008, Atkin *et al*., 2005, Tjoelker *et al*., 2001). Our results show that temperature acclimation tends to maintain homeostasis of tissue respiration below the optimum temperature for growth, indicating the importance of including temperature acclimation of plant respiration in PBMs. Studies that incorporate leaf respiratory temperature acclimation in models demonstrate an improved model capacity to reproduce observed net ecosystem exchange of CO_2_ (Smith et al., 2016), and a reduced long-term impact of global warming on vegetation carbon balance (Huntingford et al., 2017).

Our data also show a significant acclimation of photosynthetic biochemistry to growth temperature, with large changes in *V*_cmax_ and *J*_max_, and adjustments in their temperature responses resulting in an increase in temperature optimum. The changes in temperature optimum for photosynthesis due to the temperature acclimation of photosynthetic biochemistry has been well discussed and quantified in previous literature (Kattge & Knorr, 2007, Smith & Dukes, 2013, Way & Sage, 2008, Kumarathunge et al., 2018), and its implication for modelling the global C balance has been tested (Chen & Zhuang, 2013, Lombardozzi et al., 2015, Smith et al., 2016, Smith et al., 2017). As found in this paper, changes in optimum temperature of photosynthesis do not show a sizable impact on final biomass at growth temperatures below the optimum, but the effect is positive at higher temperatures (Cheesman & Winter, 2013; Way & Sage, 2008). Hence, acclimation of the optimum temperature for photosynthesis is likely to be most important when modelling responses at higher growth temperatures.

The observed decrease in *V*_cmax_ and *J*_max_ measured at 25 °C with increasing growth temperature had a large negative impact on growth. Reductions in these parameters with growth temperature have also been reported elsewhere (Way & Oren, 2010; Lin et al., 2013; Ali et al., 2015; Scafaro et al., 2017; Crous et al., 2018; Smith & Dukes, 2018), although these reductions are not consistent across the literature. In their global scale meta-analysis, Kumarathunge et al., (2018) found no overall effect of growth temperature on *V*_cmax25_ but they reported a decrease in *J*_max25_ of mature plants growing in their native environments. However, Kattge & Knorr (2007) reported no significant effect of growth temperature on either *V*_cmax25_ or *J*_max25_. A deeper understanding of how these parameters shift with growth temperature is required to parameterise this response. However, the decrease in *J*_max25_:*V*_cmax25_ from low to optimum growth temperatures is very consistent with past studies and can readily be incorporated in models (see Kumarathunge et al., 2018, Smith et al., 2020, Kattge & Knorr., 2007).

We also observed other changes in leaf gas exchange parameters with temperature. The stomatal conductance parameter, *g_1_* showed an increasing trend with growth temperature, consistent with some theoretical predictions (Prentice et al. 2014) and in common with other studies (Wang et al., 2017a; Wang et al., 2017b, Lin et al., 2015). In our study, this increase in *g*_1_ had relatively little effect on growth, compared to the changes in photosynthetic biochemistry. However, stomatal conductance values were > 0.5 mol m^-2^ s^-1^ at all growth temperature treatments, reflecting the well-watered conditions under which the seedlings were grown. It seems likely that stomatal conductance of these seedlings was not low enough to have an impact on net photosynthesis or growth. In plants with lower stomatal conductance, an increase in *g*_1_ with temperature can have a positive impact on growth.

Estimated effects on growth of changes in parameters of the photosynthetic light response curve, namely the quantum yield of electron transport (*α_J_*) and the curvature of the irradiance response curve (*θ*), were modest at both sub and supra optimal growth temperatures. Both *α*_J_ and *θ* increased from low to optimum temperature growth temperatures, and therefore had a positive impact on growth. However, at supra optimal growth temperatures, both parameters decreased, with a negative impact on growth. Reductions in *α_J_* and *θ* have previously been reported in response to low temperature stress (e.g., Groom & Baker, 1992; Rogers et al., 2019) but there are few data available to confirm our observations at high growth temperature. Many PBMs currently assume that these parameters are constant for all plant functional types and insensitive to growth temperature (Dietze, 2014; Rogers et al., 2017). Collectively, these results suggest that accounting for temperature-dependent reductions in *α_J_* and *θ* in PBMs could be important for robust simulation of plant growth as the climate warms, but that more experimental evidence is required to understand if there are general patterns in the response to warming (Rogers et al., 2019).

### Effects of warming on plant growth and allocation

Our results strongly suggested that a non-structural carbohydrate storage pool was necessary to adequately simulate growth, even in these small seedlings. Our results are consistent with several other empirical and modelling studies which have demonstrated the importance of storage C pools in modelling tree growth (Richardson *et al*., 2010; Campany *et al*., 2017; Mahmud *et al*., 2018; Fatichi et al., 2019). There have been some attempts to provide a framework to incorporate NSC storage and utilisation in PBMs (De Kauwe et al., 2014, Jones et al., 2019). However, the scarcity of data on how carbohydrate utilization and C partitioning to storage vary in response to environmental stress makes it challenging to represent these processes in PBMs (Fatichi et al., 2019). Our results suggest a significant effect of temperature on the carbohydrate utilization rate, *k*. Given that many process rates increase with warming, we had anticipated an increase in *k* with warming, similar to Jones et al. (2019) who assumed an exponential effect of temperature on the NSC utilisation rate. Instead, our analysis estimated that *k* followed a peaked temperature response, where the highest *k* was observed below the optimum temperature for other processes. However, we did not have measurements of non-structural carbohydrates available to verify our inference about the utilisation rate (Mahmud et al. 2018).

Hence, additional experiments quantifying rates of NSC accumulation and utilisation at varying growth temperatures would be valuable to develop better parameterisations for inclusion in models.

Our results also demonstrate that the allocation fractions among different tissues change in response to warming. We found that warming increased C allocation to wood, and decreased allocation to roots, but did not change foliar allocation. These observations were consistent with several empirical studies that suggested warming increases allocation aboveground at the cost of allocation belowground. For example, Blessing et al., (2015) and Rubio et al. (2025) provide evidence for decreased carbon allocation into the roots compared to leaves in *Fagus sylvatica* saplings. Similar responses have been reported for several field grown forest tree species (Drake *et al*., 2019a, Drake et al., 2019b; Melillo *et al*., 2011, Poorter *et al*., 2012). With increasing temperature, increasing plant respiration may push plants to allocate more C to non-structural C pools than structural growth; therefore, the allocation patterns may periodically change with warming (Merganicová et al., 2019, Collalti et al., 2018). Additionally, warming accelerates litter decomposition rates releasing more nutrients from soil nitrogen pool which ultimately leads to increase partitioning to aboveground tree biomass (Poorter et al., 2012; Melillo et al., 2011; Litton & Giardina, 2008). Hence, our data agree with other studies that suggest incorporating a dynamic response of C allocation in PBMs is crucial for the simulations of the impact of warming on plant growth (Merganicová et al., 2019). However, a deeper process understanding is required to parameterise this response to warming.

Our results also showed a strong effect of SLA on growth, notably at sub-optimal growth temperatures. The observed increase in SLA with growth temperature in our study is comparable with similar past studies with seedlings for example, *Eucalyptus* spp. (Drake et al 2017, Drake et al., 2015), Boreal tree species (Tjoelker et al., 1998), and several other herbaceous and woody plant species (Rubio et al. 2025, Poorter et al., 2012, Poorter et al. 2010, Poorter et al. 2009, Scheepens et al., 2010, Atkin et al., 2009). Rosbakh et al., (2015) also showed evidence for positive SLA–temperature relationship for species occurring in calcareous grasslands along a temperature gradient in the Bavarian Alps. However, our results contrast with several other studies which reported either a decrease or no change in SLA in response to warming (Bruhn et al., 2000; Way & Sage 2008; Carter et al., 1997). An increase in SLA at sub-optimal growth temperatures presumably increases total plant light interception, resulting in increased C uptake and growth (Ghannoum et al., 2010). However, at supra-optimal growth temperatures, warming reduces total leaf area leading to reduced light interception and growth (Way & Sage 2008).

Many PBMs assume SLA as a constant, but our results provide evidence for the use of temperature-dependent SLA in models for improved predictions.

The modelled rate for C losses through growth respiration, root exudates and volatile organic C emission, *Y,* showed a peaked response with increasing temperature. The lower rates of *Y* observed at higher growth temperatures suggest that the extra C cost for biomass production is lower at high temperatures which may potentially reflect a structural change in biomass composition such as changes in chemical composition (Villar & Merino, 2001; Poorter, 1994). Also, the higher rates of *Y* observed at growth temperatures around the optimum temperature for growth would suggest that respiration is up-regulated when labile C accumulates, an assumption embedded in several PBMs (Zaehle & Friend, 2010). However, in this experiment, we inferred the growth respiration rates from the DA framework; therefore, the values could possibly also include C losses via other pathways such as root exudates and volatile organic C emission (Emmerson et al., 2020). Direct measurement of these C loss pathways and their interactions with environmental variables such as temperature is difficult. As a result, incorporation of these responses in PBMs remains difficult. Nevertheless, the DA approach provides a way of understanding what *Y* would have to be if the measured rates of photosynthesis, respiration and growth were to be consistent, and therefore provides a framework for further studies in the environmental control of growth respiration and other C loss pathways.

### Broader applicability of the analysis

A significant advantage of the approach used here is that it provides a way to link the observed responses to the underlying measured processes and thus develop a more mechanistic explanation of the temperature response of tree growth. Classical growth analysis is often used to interpret results of seedling experiments but does not readily allow tree growth responses to be linked to underlying observations of tree physiological responses in a manner that can be directly translated to PBMs. Drake *et al*. (2017) applied classical growth analysis to the same experiment as used here. They observed that net assimilation rate (NAR) tended to decline with growth temperature, but the temperature dependence of *Eucalyptus* seedling growth was determined primarily by leaf size and specific leaf area. Although this approach can provide insight into temperature effects on growth, it leaves some important questions unanswered, such as why the net assimilation rate declines with increasing temperature despite the strong peaked temperature response of net photosynthetic rate (see Pooter, 2002; Poorter and Navas, 2003; Tjoelker et al. 1999; Tjoelker et al. 1998a; Tjoelker et al. 1998b).

In contrast, the DA framework used in this study connects these disparate experimental measurements and thus provides new insights into determinants of plant growth. The major challenge of the approach is ensuring that the range of measurements made is sufficient to constrain the model. The full suite of measurements required includes photosynthesis, respiration, biomass growth over time, and non-structural C concentrations. However, other studies using DA approaches have been able to constrain model parameters with a less extensive set of data (Au *et al*., 2023, Williams et al., 2005).

It would be valuable to apply this approach to a broader range of studies to obtain a more general picture about the mechanisms that determine the temperature response of tree growth in trees, including studies in cooler and warmer environments, and studies on larger trees. We note that the experimental data considered here came from seedlings that were well-watered throughout the experimental period. Water limitation reduces the temperature optima for both photosynthesis and growth, total leaf area display and specific leaf area (Kumarathunge et al., 2020). The analysis presented here does not consider the constraints placed on temperature responses by water availability (Allen et al., 2010, Barber et al., 2000, Densmore-McCulloch et al., 2016), but it could potentially be extended to examine the indirect effects of warming through increased water limitation.

## Conclusion

Using a DA framework, we quantified key mechanisms that contribute to the temperature response of tree seedling growth. We show that not only photosynthesis and respiration, but also other C balance processes including non-structural carbohydrate utilisation rate, growth respiration and C allocation to different tissues are also important in determining tree growth responses to increasing temperature. The DA framework used in this study provides an approach to connect disparate experimental measurements and provides new insights into determinants of plant growth. This approach is useful to obtain a more holistic understanding of the C balance in plants from experiments which ultimately should prove useful in quantifying temperature limits to tree growth and improving the quantification of tree growth by PBMs.

## Data Sharing and Data Accessibility

Part of the dataset used for this study is publicly available (Drake et al., 2017) and the rest of data and analysis code to reproduce all the results, including the figures and tables, is available as a *git* repository (https://bitbucket.org/Kumarathunge/greatda).

## Acknowledgements

This research was supported by Australian Research Council Discovery Projects DP140103415 and DP160103436 with additional support from the Hawkesbury Institute for the Environment and Western Sydney University. DK was supported by a Western Sydney University international PhD scholarship. We thank Gavin Mckenzie, Goran Lopaticki and Burhan Amiji for technical assistance.

## Author contribution statement

Project conceived by BEM. Data collection and analyses designed and carried out by DPK and KM with guidance from BEM. FJC contributed to data collection. Manuscript writing led by DK with contributions from BM and KM. JED and MGT contributed to data collection and made a substantial contribution to the experimental design, data interpretation, and writing. DK and KM equally contributed to this paper.

## Methods S1

### Experimental design, growth and physiological data measurement and parameterising photosynthesis and respiration

This experiment was carried out at the Hawkesbury Institute for the Environment, Western Sydney University, Richmond NSW, Australia (33°37’S 150°44’E). Seedlings of three provenances (tropical, sub-tropical and temperate) of forest red gum (Eucalyptus tereticornis sp tereticornis) were transplanted in to 7-L PVC pots filled with a moderately fertile sandy loam soil. Data were pooled across provenances in this analysis because a separate analysis using data from this experiment demonstrated that widely distributed E. tereticornis provenances share a common physiological thermal niche without local adaptation to the climate of origin (Drake et al., 2017b). We randomly placed 30 seedlings in each of six adjacent naturally sunlit glasshouse rooms in the austral summer on 2016-01-08 (day 0). We implemented six daily mean growth temperature treatments, which averaged 18, 21.5, 25, 28.5, 32 and 35.5 °C. Each temperature treatment consisted of 10 temperature set points with a 9 °C diurnal range around the mean temperature values during the day-night cycle. Seedlings were kept well-watered using an automated irrigation system and were fertilized fortnightly with a liquid fertilizer (Aquasol, Yates Australia; 250 ml per seedling). We recorded air temperature, VPD, RH and photosynthetically active radiation (PAR) in one-minute intervals day and night and monitored conditions frequently to maintain the desired control levels. Relative humidity was kept above 50%, but vapour pressure deficit (VPD) increased with growth temperature (∼0.5 kPa at 18 °C and ∼2 kPa at 35.5 °C). A detailed description of the experimental design can be found in Drake et al. (2017).

We measured height of the main stem and basal diameter of 15 seedlings per growth temperature at approximately weekly intervals. We used growth-temperature specific allometric models to estimate total leaf area and total C in leaf, wood and root pools (*C*_t,f_, *C*_t,w_ and *C*_t,r_ respectively) on a weekly time-step from the measured stem height (h) and diameter (d). The allometric models were developed using seedlings that were periodically harvested throughout the study (*n*=156). The allometric models were of the form:

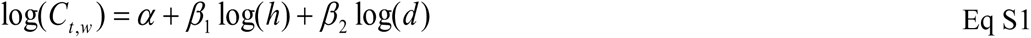

The allometric models predicted the observed leaf area and component masses with high accuracy (*r*^2^ > 0.95 for all growth temperature treatments; Drake et al. 2017b).

We measured leaf net photosynthetic rate (*A*_net_) midway through the experiment (26 days after planting) and prior to the final harvest (44 days after planting). Measurements were made on new fully expanded leaves on eight replicate seedlings in each growth temperature treatment. We used six Licor 6400XT portable photosynthesis systems (Licor Biosciences, Lincoln, NE, USA), with the standard 2×3 cm leaf chamber and LED light source (Li-6400-02B LED).

Measurements were conducted at four different photosynthetically active radiation levels (PAR); 1500, 1000, 500 and 100 µmol m^-2^ s^-1^. The relative humidity inside the leaf chamber was maintained between 60 – 80%. The sample CO_2_ concentration was held at 400±5 ppm and flow rate at 500 µmol s^-1^. We measured *A*_net_ at a leaf temperature similar to the mid-morning growth temperature (20, 24, 28, 32, 36, and 40 °C respectively). We also measured the direct short-term temperature response of light-saturated photosynthesis (*A*_sat_) on day 29 after planting using seedlings from the 21.5 °C growth temperature treatment only (*n* = 8). Seedlings were moved among treatment rooms and measured at all six temperatures. A minimum wait period of 20 min allowed the seedlings to adjust to the new temperature before measurement. These data were published, in part, in Drake *et al*., (2017b).

We measured photosynthetic CO_2_ response curves (A/C_i_ curves) on a random subset of seedlings (*n* = 6) of three growth temperature treatments (18, 28.5 and 35.5 °C) prior to final harvest (40 – 44 days after planting). Measurements were started at ambient CO_2_ level (400 µmol mol^-1^) and changed stepwise through a series of sub-ambient (40-400 µmol mol^-1^) to super-ambient saturating CO_2_ concentrations (400-2000 µmol mol^-1^). We repeated the same measurement protocol on the same leaf at five different leaf temperatures.

We measured the short-term temperature response of leaf dark respiration at night (22:00-02:00 h), 35 or 36 days after planting, using seedlings from the 21.5 °C growth temperature treatment only. These measurements were conducted at five night-time temperatures (14, 18, 22, 26, and 30 °C) using five Li-6400-XT machines equipped with conifer chambers (Li-6400-22; Licor). Measurements were made on single detached leaves of eight plants (*n*=8). Leaves were placed in moist polythene bags and moved among treatment rooms for measurement. Each leaf was allowed to adjust to each growth temperature for at least 20 min in a darkened box before measurements commenced. These data were published in Drake *et al*., (2017b).

We also measured leaf, stem and root respiration rates for five seedlings at each growth temperature (i.e. acclimated temperature response) during the final harvest (44-48 days after planting). We measured leaf respiration rates using three randomly selected leaves, combined in a single cuvette for measurement, for each of the replicate seedlings. We used the entire stem with branches cut into 5 cm segments to measure stem respiration rates. We used entire root system or subsample, depending on root mass, to measure root respiration rates. All tissue-specific respiration rates were measured at a common temperature of 25 ± 1.5 °C and a reference cell CO_2_ concentration of 400±5 ppm, using Licor 6400XT portable photosynthesis systems with the Li-6400-22 conifer chamber. A detailed description of this measurement protocol and resulting data can be found in (Drake *et al*., 2017b).

### Parameterising photosynthesis and respiration

Following Campany *et al*., (2017), we used a coupled photosynthesis-stomatal conductance model to quantify the leaf net photosynthetic rate of seedlings at each growth temperature. The standard biochemical model of photosynthesis (Farquhar *et al*., 1980) was coupled to the optimal stomatal conductance model proposed by Medlyn *et al*. (2011a). The model was run with photosynthetic parameters and meteorological data measured at each growth temperature treatment. We quantified the photosynthetic parameters using the gas exchange data as described below.

We estimated plant photosynthetic capacity by fitting the A/Ci curves using the Farquhar *et al*. (1980) (FvCB) biochemical model of photosynthesis. We used the *fitacis* function within the *plantecophys* R package (Duursma 2015) in R version 3.5.1 (R Development Core Team, 2018) to estimate the key parameters of the FvCB model: the maximum rate of ribulose-1,5-bisphosphate carboxylase-oxygenase (Rubisco) activity (*V*_cmax_) and the potential rate of electron transport (*J*_max_). We assumed the Bernacchi *et al*. (2001) kinetic constants for the temperature response of Michaelis–Menten coefficients of Rubisco activity and CO_2_ compensation point in the absence of mitochondrial respiration.

We used the leaf net photosynthesis measurements measured at in-situ growth temperatures to estimate the stomatal conductance model parameter (*g_1_*) following Medlyn *et al*. (2011). We quantified the temperature response parameters of *V*_cmax_ and *J*_max_ following Kumarathunge *et al*. (2018b). As the temperature response parameters of *V*_cmax_ and *J*_max_; *E*_a_ and *ΔS* were estimated only for three growth temperatures (18, 28.5 and 35.5 °C), we assumed the same *E*_a_ and *ΔS* for growth temperatures 18 and 21.5°C, 25 and 28.5°C, 32 and 35.5 °C respectively. We used non-linear regression to simultaneously estimate the parameters of the coupled photosynthesis-stomatal conductance model, namely *V*_cmax_ and *J*_max_ at standard temperature of 25 °C, quantum yield of electron transport (*αj*) and the curvature of the light response of electron transport rate (*θ*), from *A*_net_ measurements at different PAR levels at two time points (26 and 48 days after planting) for all growth temperature treatments. Parameter estimation used the fitted values of *g_1_*, *E*_a_*, ΔS* and measurement leaf temperature, VPD and CO_2_ concentration as inputs. This approach optimised the model parameters against the measured *A*_net_ data.

Similar to the measured *A*_sat_ values, optimized *V*_cmax_ and *J*_max_ values showed a clear decline between 26 and 48 days after planting at growth temperatures above 18 °C. However, at 18 °C, both parameters increased. Therefore, we assumed that both *V*_cmax_ and *J*_max_ (at a standard temperature of 25 °C), linearly decrease between 26 and 48 days after planting at growth temperatures above 18 °C (Fig. S3). We have implemented these temporal variations of both *V*_cmax_ and *J*_max_ in estimating GPP at different growth temperatures.

## Notes S1

### Short-term and acclimatory effects of temperature on physiological processes

The short-term temperature dependence of leaf-level photosynthesis at saturating light (*A*_sat_) (i.e., *A*_sat_ measured at varying leaf temperatures on plants grown at 21.5 °C) showed a peaked relationship with leaf temperature (Fig. S1a). The optimum temperature for *A*_sat_ (*T*_optA_) was 29.1 (±0.23) °C. Compared to the short-term temperature response, the acclimated temperature response (i.e., *A*_sat_ measured at leaf temperature similar to the mid-morning growth temperature on plants grown at each growth temperature) showed a slightly lower *T*_optA_ (27.7 °C). The *A*_sat_ at the optimum was higher for the short-term response (27.2 µmol m^-2^ s^-1^) compared to the acclimated response (23.1 µmol m^-2^ s^-1^).

Leaf dark respiration showed an exponential increase with short-term changes in leaf temperature, with a *Q*_10_ of 2.1 (solid line, Fig. S1b). Leaf dark respiration rates measured at growth temperatures (dashed line, Fig. S1b) showed clear acclimation to temperature. Rather than increasing exponentially with temperature, the tissue-specific respiration rates did not vary between 18 and 25 °C. At growth temperatures above 25 °C, both leaf and root respiration rates increased. Leaf respiration rate increased markedly, indicating a lack of homeostasis (Fig. S1b). Notably, acclimation of wood respiration resulted in homeostasis across the full range of growth temperatures (Fig. S4a). Similar to leaves, the long-term temperature response of root dark respiration rate showed a homeostatic acclimation at low to mid growth temperatures (18-25 °C; Fig. S4b).

The instantaneous temperature responses of both *V*_cmax_ and *J*_max_ showed a peaked pattern with leaf temperature (Fig. S1c, S1d). The short-term temperature responses of both *V*_cmax_ and *J*_max_ clearly diverged between growth temperatures, indicating longer-term temperature acclimation. The optimum temperatures of both *V*_cmax_ and *J*_max_ were increased with increasing growth temperature. The optimal temperature at the highest growth temperature was 1.5 °C higher than the optimum temperature at the lowest growth temperature. For *J*_max_, this difference was 2.2 °C. The rates of both *V*_cmax_ and *J*_max_ measured at 25 °C decreased significantly with increasing growth temperatures (Fig. S1c, S1d). However, the rate of decrease in *J*_max_ was higher than the *V*_cmax_ at growth temperatures below 28.5 °C, therefore we observed a decrease in *J*_max_:*V*_cmax_ from growth temperatures 18 °C (1.8) to 28.5 °C (1.4). The ratio was similar between growth temperatures 28.5 °C and 35.5 °C. Further, the activation energies of both *V*_cmax_ and *J*_max_ were lower at low growth temperatures compared to the values observed at highest growth temperature (Table S1).

Other physiological parameters also showed acclimation to growth temperature (Table S1). The stomatal conductance parameter (*g*_1_) showed an increasing trend with growth temperature (Table S1). The lowest *g*_1_ value (5.5 kPa^0.5^) was observed at 18 °C and the highest (29.0 kPa^0.5^) was observed at 35.5 °C. We observed a peaked response for the quantum yield of electron transport with the highest value observed at 28.5 °C (Table S1). Light saturated net photosynthesis rates and *V*_cmax_ and *J*_max_ (at a standard temperature of 25 °C) declined between the two measurement times (26 and 48 days after planting) at growth temperatures above 18 °C (Fig. S2, Fig. S3). At 18 °C, *A*_sat_ increased with time. We speculate that the decline in net photosynthesis, *V*_cmax_ and *J*_max_ at growth temperatures above 18 °C could be mainly due to the changes in seedling growth rates as influenced by external factors such as sink limitation in these fast-growing seedlings (Campany et al., 2017). Together, these results demonstrate the importance of both short-term and long-term temperature responses on photosynthetic and respiratory physiology. We incorporated these dynamic changes in photosynthetic rates when estimating the daily GPP through dynamically changing *V*_cmax_ and *J*_max_ (as shown in Fig. S3) in the coupled photosynthesis-stomatal conductance model.

**Fig. S1.**
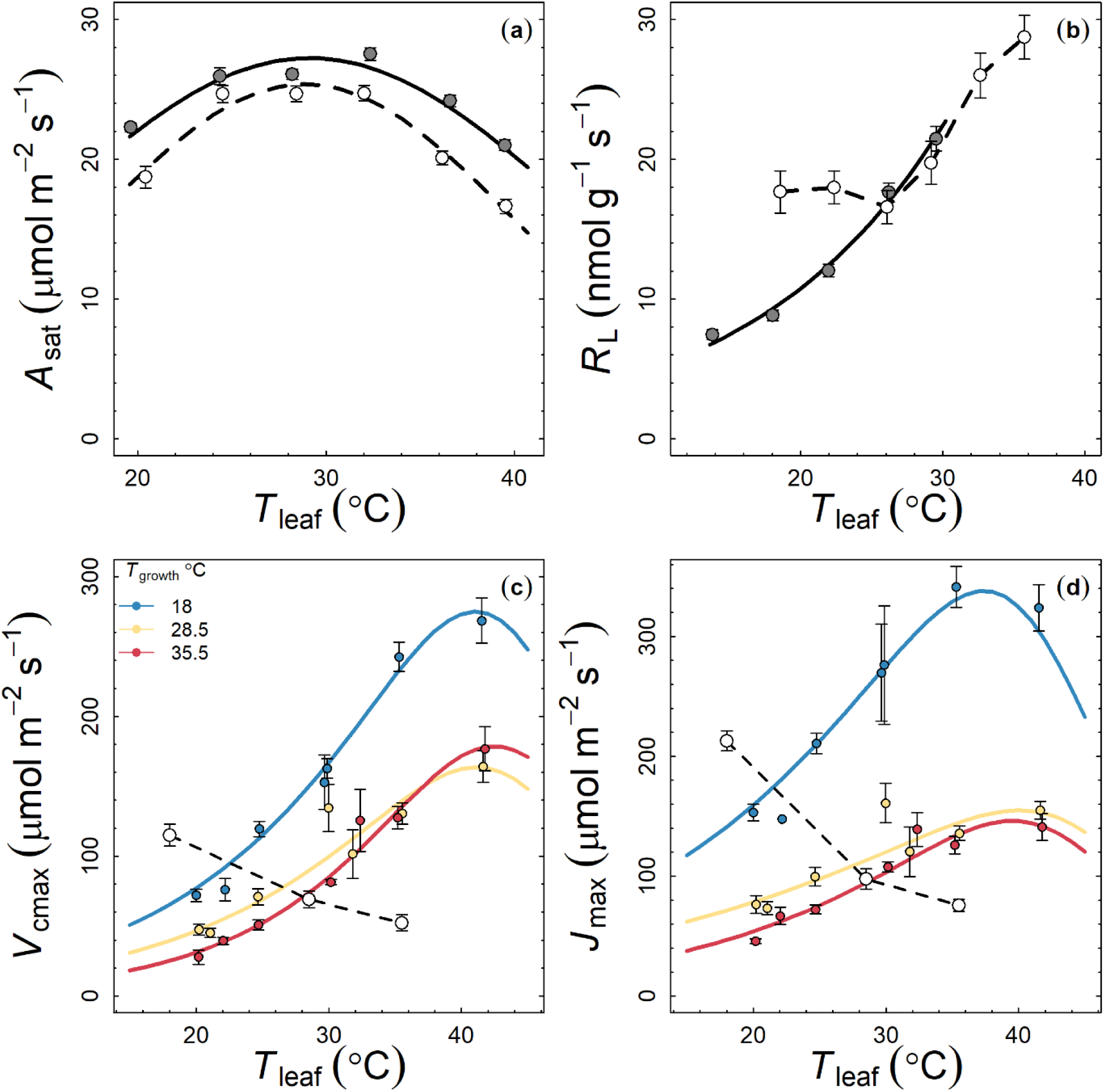
The temperature response of leaf photosynthesis (a), leaf dark respiration (b) maximum rate of ribulose-1,5-bisphosphate carboxylase-oxygenase (Rubisco) activity (*V*_cmax_, c) and potential rate of electron transport (*J*_max_, d) of *E. tereticornis*. In panels (a), filled circles and continuous lines depict the direct short-term temperature responses (i.e. *A*_sat_ measured at varying leaf temperatures on plants grown at 21.5 °C) and empty circles with dashed lines depict the acclimated temperature responses (where measurement were taken at a leaf temperature similar to the mid-morning growth temperature (20, 24, 28, 32, 36, and 40 °C respectively on plants grown at each growth temperature) of leaf photosynthesis. In panel (b), filled circles and continuous lines depict the direct short-term temperature responses and empty circles with dashed lines depict the acclimated temperature responses (rates were measured at a common temperature of 25 ± 1.5 °C at each growth temperature) of leaf respiration. In panels (c) and (d), continuous lines in different colours depict the direct short term temperature response of *V*_cmax_ and *J*_max_ measured at three different growth temperatures and the empty circles with dashed lines depict the acclimated temperature responses (i.e. *V*_cmax_ and *J*_max_ values at a common temperature of 25 °C). Error bars (where visible) depict ±1SE (n = 8 for photosynthesis and n=5 for leaf dark respiration, *V*_cmax_ and *J*_max_).

**Fig. S2.**
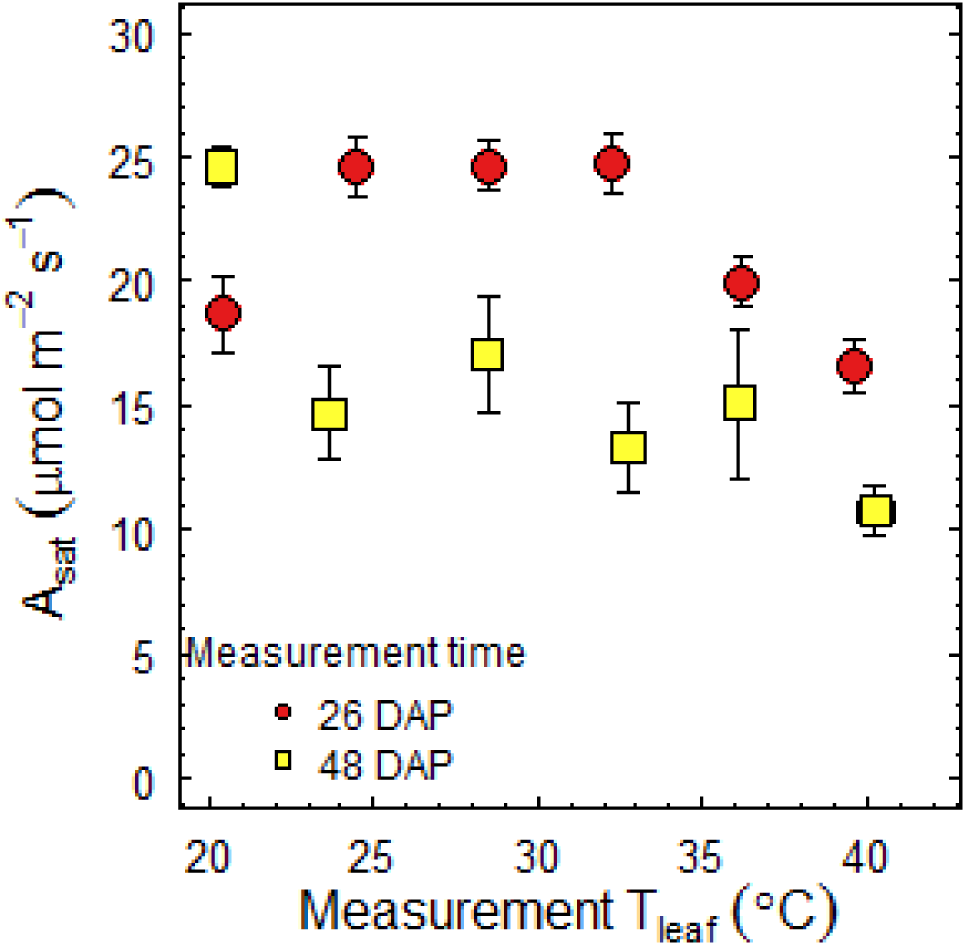
Irradiance saturated net photosynthetic rates (A_sat_) at in-situ growth temperatures measured at two different time points; 26 and 48 days after planting (DAP).

**Fig. S3.**
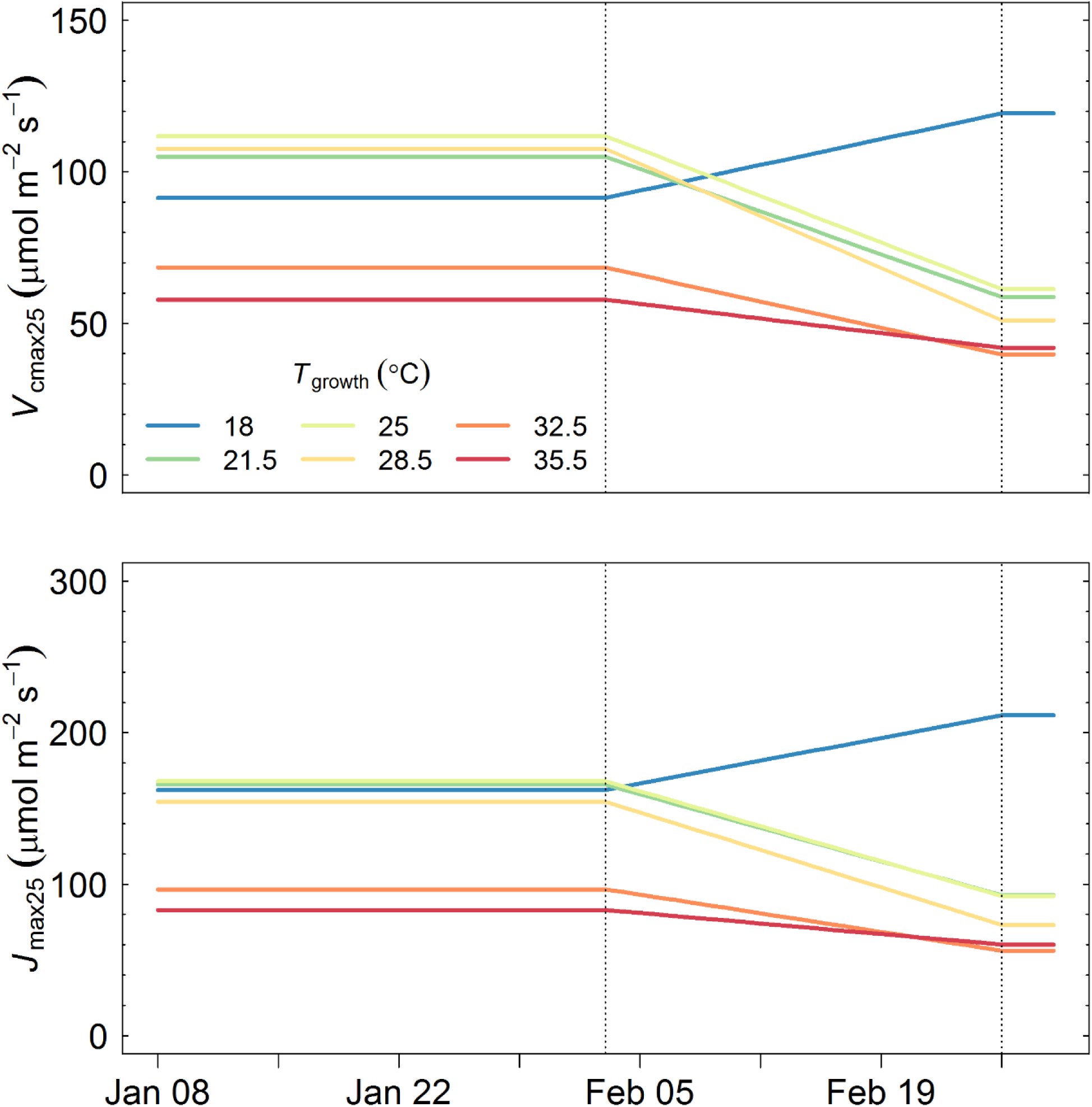
Temporal variation of *V*_cmax_ and *J*_max_ at a standard temperature of 25 °C at different growth temperature treatments.

**Fig. S4.**
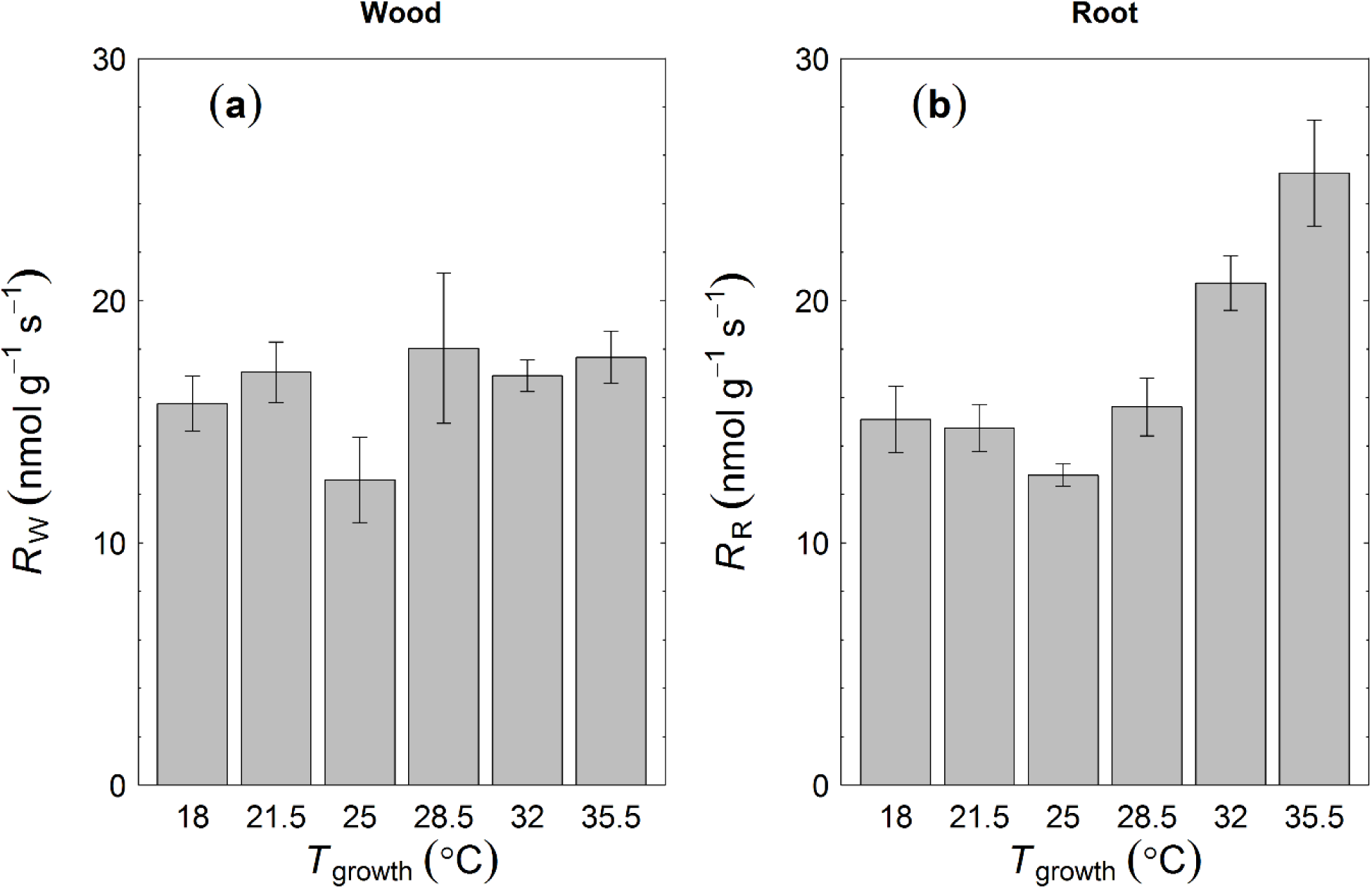
Tissue-specific respiration rates of wood (a) and roots (b) of *E. tereticornis* seedlings measured at growth temperatures. Wood and root respiration rates are expressed per unit mass. Error bars depict ±1SE.

**Fig. S5.**
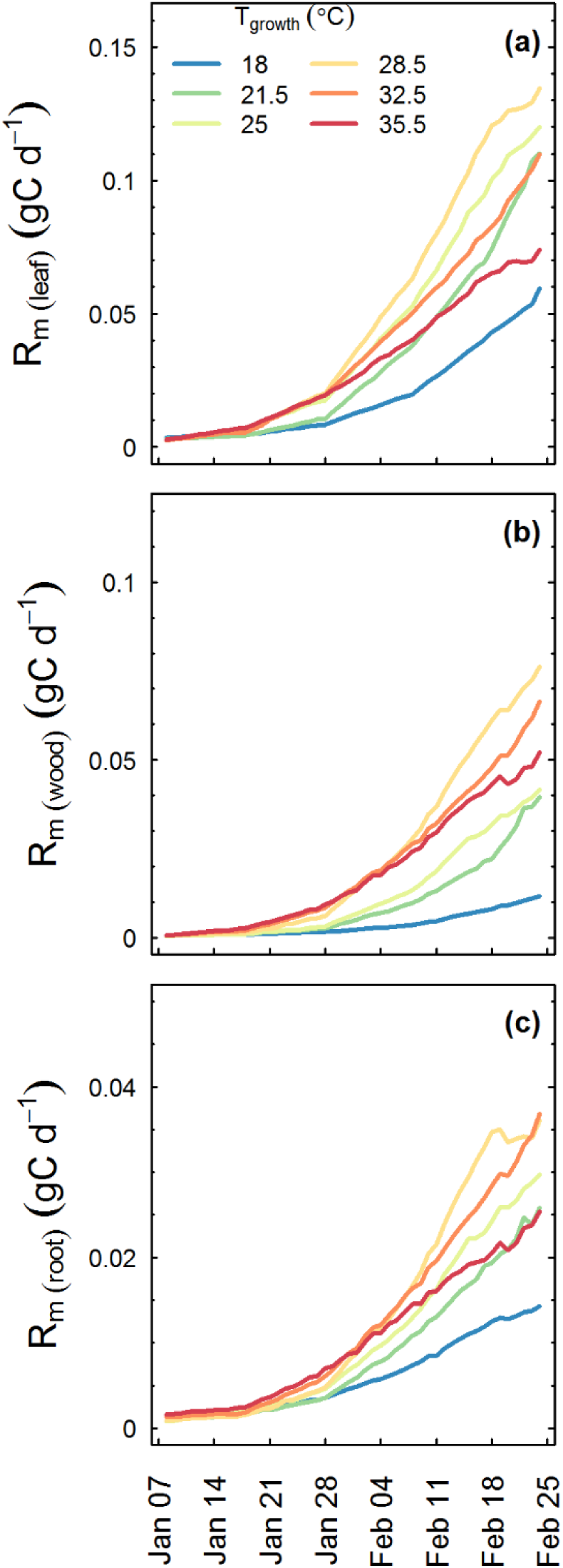
Time series of daily respiration (R_m_) of different tissues: (a) Leaves, (b) wood and (c) roots. Major ticks on the x-axis reflect weeks. Note the differences in Y axis scale in different panels.

**Fig. S6.**
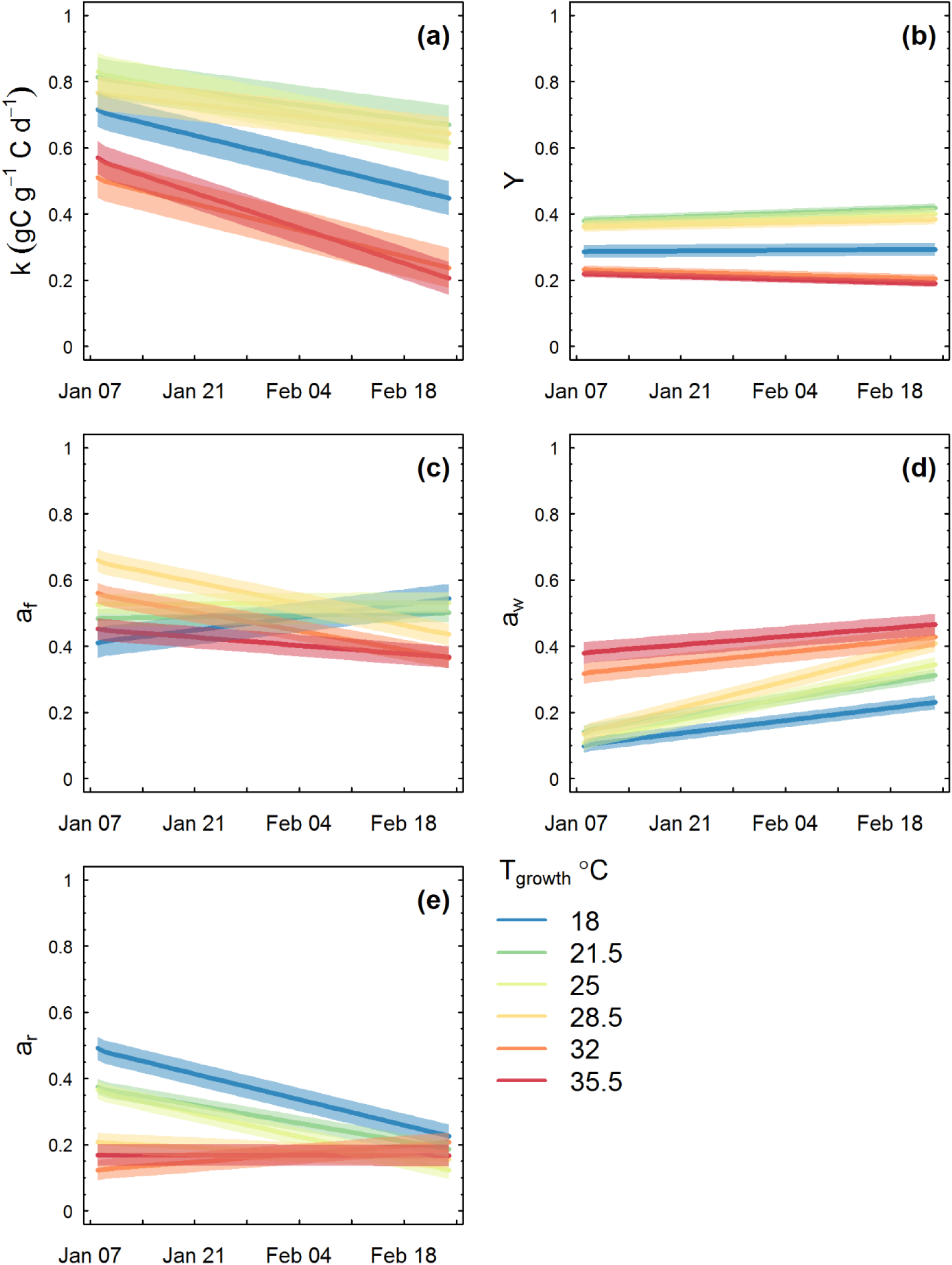
Time series of parameters of the carbon balance model: (a) storage utilization coefficient, k; (b) unmeasured C losses, Y; (c) allocation to foliage, a_f_; (d) allocation to wood, a_w_; (e) allocation to roots, a_r_. Lines depicts the fitted linear regression model with the shaded area showing the 95% CI of predictions.

**Fig. S7.**
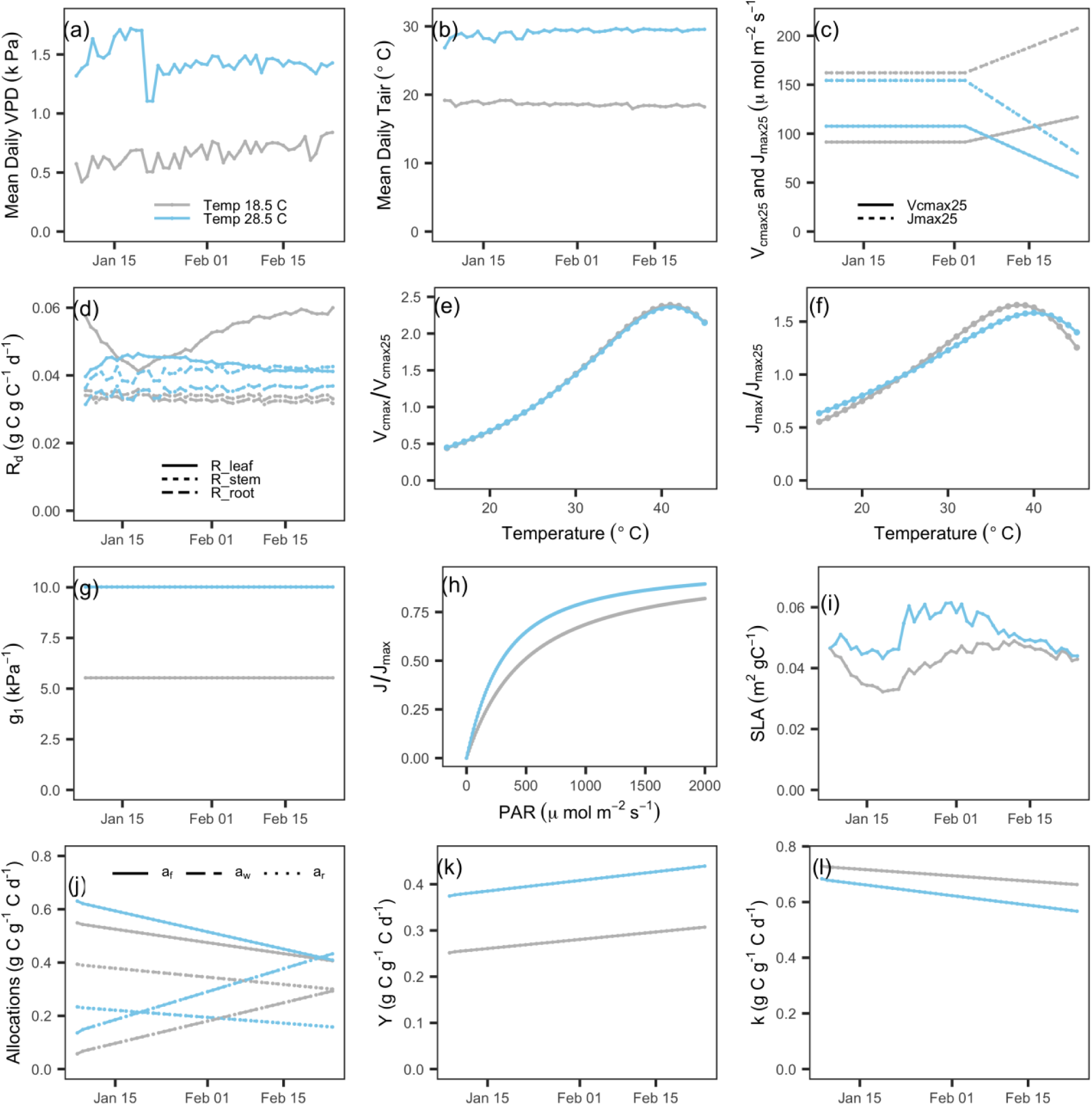
Changes in input parameters of the attribution analysis from sub-optimal to optimal growth temperatures.

**Fig. S8.**
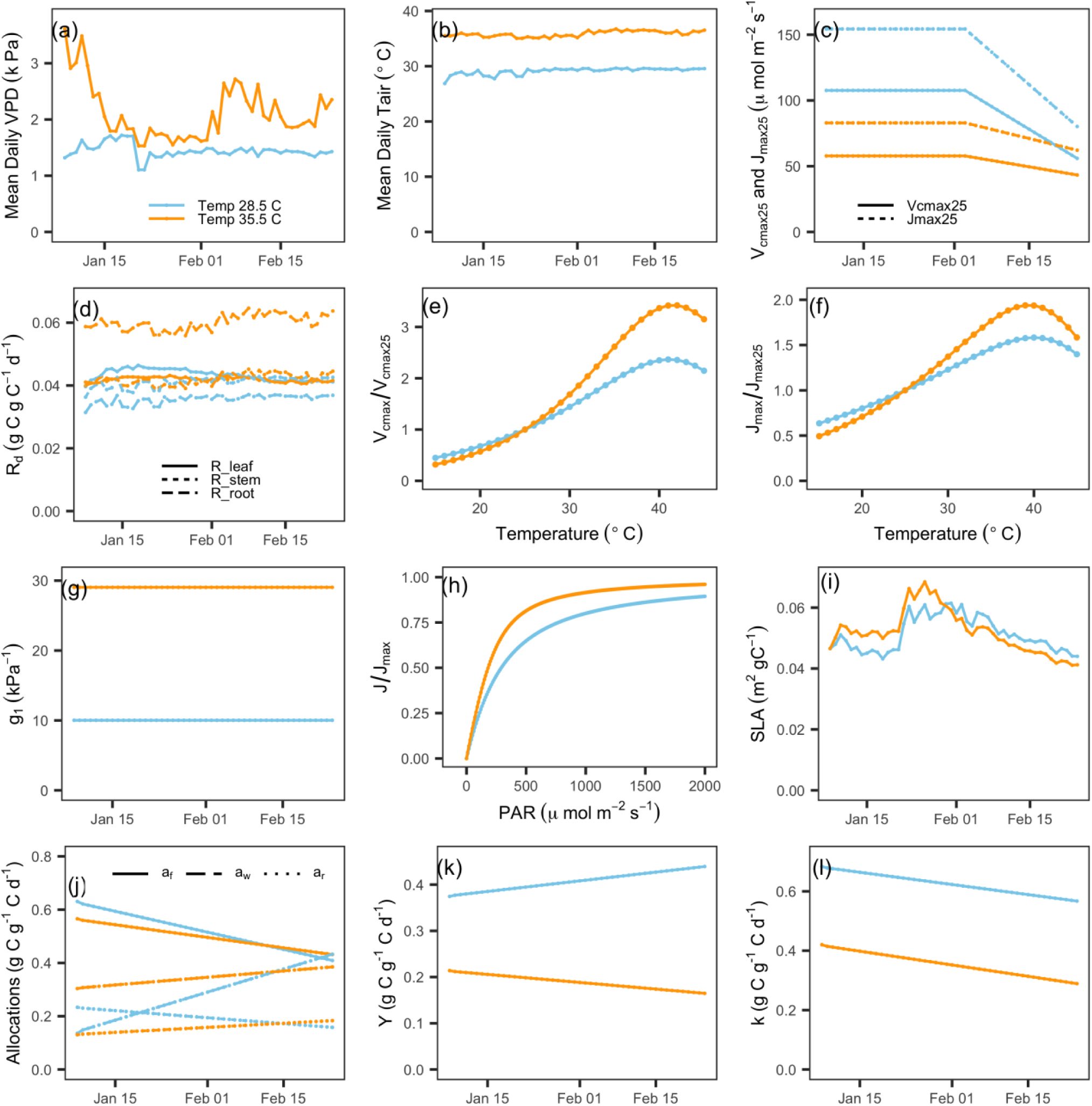
Changes in input parameters of the attribution analysis from optimal to supra optimal growth temperatures.

**Table S1.**
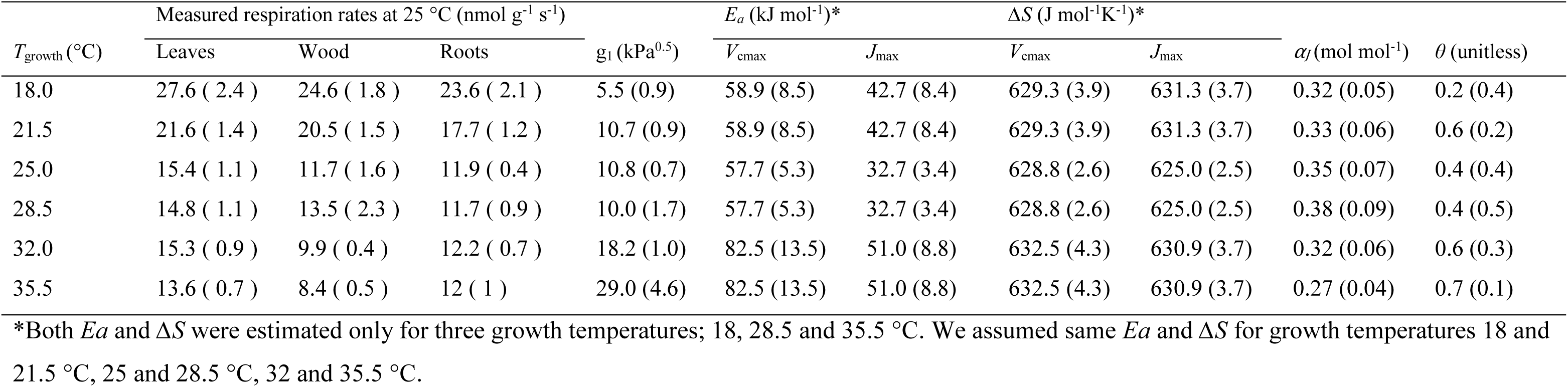
Tissue specific respiration rates at a common temperature of 25 °C at different growth temperatures that were used to estimate total seedling respiration and physiological parameters that were used to estimate daily photosynthetic rate per unit leaf area (GPP; gC m-2 d-1) at different growth temperatures using the coupled photosynthesis-stomatal conductance model.

